# Re-evaluating the role of nucleosomal bivalency in early development

**DOI:** 10.1101/2021.09.09.458948

**Authors:** Rohan N. Shah, Adrian T. Grzybowski, Jimmy Elias, Zhonglei Chen, Takamitsu Hattori, Carolin C. Lechner, Peter W. Lewis, Shohei Koide, Beat Fierz, Alexander J. Ruthenburg

## Abstract

Nucleosomes, composed of DNA and histone proteins, represent the fundamental repeating unit of the eukaryotic genome^1^; posttranslational modifications of these histone proteins influence the activity of the associated genomic regions to regulate cell identity^2–4^. Traditionally, trimethylation of histone 3-lysine 4 (H3K4me3) is associated with transcriptional initiation^5–10^, whereas trimethylation of H3K27 (H3K27me3) is considered transcriptionally repressive^11–15^. The apparent juxtaposition of these opposing marks, termed “bivalent domains”^16–18^, was proposed to specifically demarcate of small set transcriptionally-poised lineage-commitment genes that resolve to one constituent modification through differentiation, thereby determining transcriptional status^19–22^. Since then, many thousands of studies have canonized the bivalency model as a chromatin hallmark of development in many cell types. However, these conclusions are largely based on chromatin immunoprecipitations (ChIP) with significant methodological problems hampering their interpretation. Absent direct quantitative measurements, it has been difficult to evaluate the strength of the bivalency model. Here, we present reICeChIP, a calibrated sequential ChIP method to quantitatively measure H3K4me3/H3K27me3 bivalency genome-wide, addressing the limitations of prior measurements. With reICeChIP, we profile bivalency through the differentiation paradigm that first established this model^16,18^: from naïve mouse embryonic stem cells (mESCs) into neuronal progenitor cells (NPCs). Our results cast doubt on every aspect of the bivalency model; in this context, we find that bivalency is widespread, does not resolve with differentiation, and is neither sensitive nor specific for identifying poised developmental genes or gene expression status more broadly. Our findings caution against interpreting bivalent domains as specific markers of developmentally poised genes.

---

In its original conception, the bivalency model posits that the combination of H3K4me3 and H3K27me3 represents a specific regulatory marker of developmentally staged genes. Specifically, lineage commitment genes are thought to be held in a low-expression, transcriptionally “poised” state by promoter nucleosomes bearing both H3K4me3 and H3K27me3^16,18,21,22^. Upon differentiation, the bivalent domain “resolves” into a monovalent state, and the associated gene is either transcriptionally activated or terminally repressed if H3K27me3 or H3K4me3 is lost, respectively^16,18,21,22^. The elegance of this instructive model inspired a host of follow-on studies that have suggested that bivalency is important in differentiation^23–30^, embryogenesis^17,31–34^, genome architecture^22,35–38^, and oncogenesis^39–43^.

In the absence of unambiguous biochemical or functional validation^17,44,45^, these studies have largely relied upon ChIP, with the vast majority of studies defining loci with independent ChIP enrichment for H3K4me3 and H3K27me3 as bivalent domains. However, this analysis cannot distinguish whether the two modifications coexist or represent two distinctly marked subpopulations of alleles or cells. Further, because different ChIPs are normalized separately, they exist on separate relative scales and cannot be quantitatively compared without internal calibration^46–48^. As such, it is impossible to quantify the extent of bivalency at a given locus or to measure its changes through differentiation.

To address the first problem, several studies have used sequential ChIP^16,29,49–51^, measuring coexistence by using the eluent of an IP against H3K4me3 as the substrate for an IP against H3K27me3 (or vice versa). However, these experiments used antibodies of unknown specificity^16,29,49–51^, were uncalibrated, and were often undersampled^50,52^, precluding quantification of the extent of modification. Moreover, many used relatively large chromatin fragments in their pulldowns, making it difficult to determine whether modifications coexisted on one nucleosome or discretely marked neighbouring nucleosomes^16,29,50,51^. The limitations of these sequential ChIP studies preclude accurate assessment of key properties of bivalency.

Our previous work introduced internally calibrated ChIP (ICeChIP), in which barcoded nucleosome internal standards are used to measure antibody specificity and as analytical calibrants that enable computation of the histone modification density (HMD), or the proportion of nucleosomes at a given locus with the modification of interest^46–48^. By identifying regions with high H3K4me3 and H3K27me3, we indirectly identified many promoters with a nonzero amount of bivalency, including those regulating developmental and metabolic genes^46^. However, this analysis was limited; it was not sensitive for bivalency at less extensively modified loci, nor could it quantify the extent of bivalency. Here, we directly quantify this nucleosomal mark pattern by calibration of a modified sequential ChIP approach to critically evaluate the bivalency model in the differentiation system in which the foundational observations were made.

## Measuring bivalency with reICeChIP

To directly measure bivalency and evaluate its role in differentiation, we first attempted to deploy our calibrants with published sequential ChIP methods. However, when evaluated with internal standards, these methods^16,29,49^ displayed extremely low enrichment and variable specificity (Extended Data Fig. 1a), with common elution methods either failing to release most of the captured material^51^ or compromising the specificity of the second IP (Extended Data Fig. 1b-c). With such heavy losses, we became concerned that we would undersample and potentially bias the measurement of bivalent nucleosomes. We sought a method of elution from the primary IP that was both more efficient and would preserve nucleosome integrity for the second IP. To that end, we modified a recombinant biotinylated Fab (304M3-B) specific for H3K4me3^53^ with an intervening HRV 3C endoprotease cleavage site to enable quantitative elution by enzymatic cleavage under mild conditions.

We then leveraged this reagent to develop reICeChIP (Fig. 1a). The first pulldown was conducted with the cleavable α-H3K4me3 Fab from native mononucleosomes^46,54^ spiked with nucleosome internal standards. We then eluted the captured nucleosomes from streptavidin resin by cleaving the antibody with HRV 3C endoprotease^55^ and, with this eluent, conducted a second pulldown against H3K27me3 with a conventional antibody. This method eluted material from the primary pulldown more efficiently (Extended Data Fig. 1d), resulting in 1000-2500x higher enrichment of the target over the published methods (Fig. 1b, Extended Data Fig. 1a). This improvement enabled genome-wide measurement of bivalency HMD (Fig. 1c), representing the proportion of nucleosomes at a given locus modified with both H3K4me3 and H3K27me3, using the trans-bivalent nucleosome standards^56^ as the calibrant (Extended Data Fig. 1e, 2; Supplementary Note 1).

**Fig. 1.**
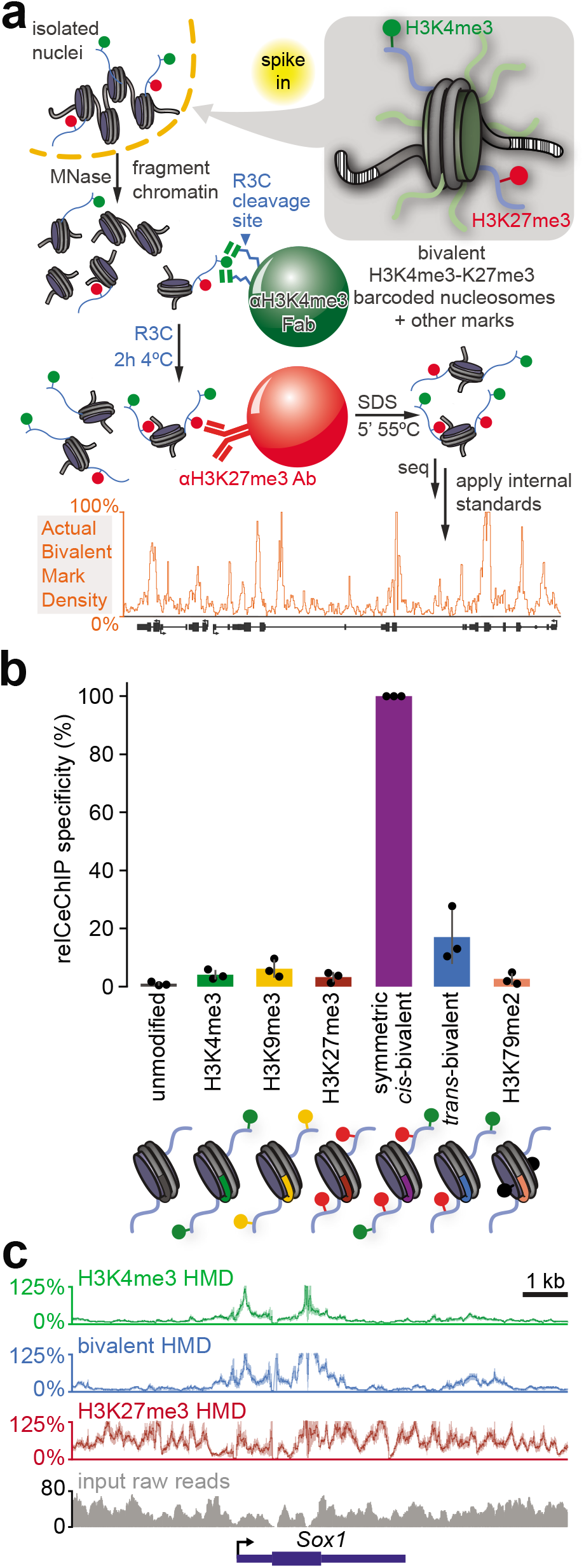
Workflow and evaluation of reICeChIP-seq. **(a)** Schematic of reICeChIP-seq. The recombinant α-H3K4me3 Fab 304M3-B achieves high affinity by “clasping” the histone tail between two Fab molecules^53^, a binding mode readily achieved by multiple copies of the Fab presented on a bead, but not by the Fab in solution. Thus, protease cleavage not only elutes nucleosomes from the beads but also likely from the Fab complex. **(b)** Enrichment of different barcoded nucleosomes in reICeChIP-seq (n=3 biological replicates). Error bars represent S.D. **(c)** Representative line plot showing histone modification density of H3K4me3, H3K27me3, and bivalency ICeChIP-seq presented with 95% confidence intervals (lighter shade) and input read depth in naïve mESCs. Bivalency is calibrated to the trans-bivalency nucleosome standard and corrected for off-target H3K9me3 pulldown.

## Bivalency through differentiation

With reICeChIP, we sought to study the role of bivalency in development by tracking its changes across a differentiation pathway that was used in several classic studies of bivalency^16,18,57^: differentiation from naïve mESCs^57^ through the primed mESC state^57^ to NPCs. In naïve mESCs, we noted that bivalency was far more widespread than previously reported (Fig. 2a-b); rather than ∼1000 bivalent genes in naïve mESCs^57^, we observed at least 10% bivalency HMD at most promoters (25768/42622), with almost 5000 promoters bearing bivalency at more than 50% of their nucleosomes (Fig. 2a,c; Supplementary Notes 2, 3). This trend is recapitulated with primed mESCs, with the consensus set of bivalent promoters representing fewer than 2000 genes^18,22,35,58^, as compared to more than 25,000 that are >25% bivalent in our analysis.

**Fig. 2.**
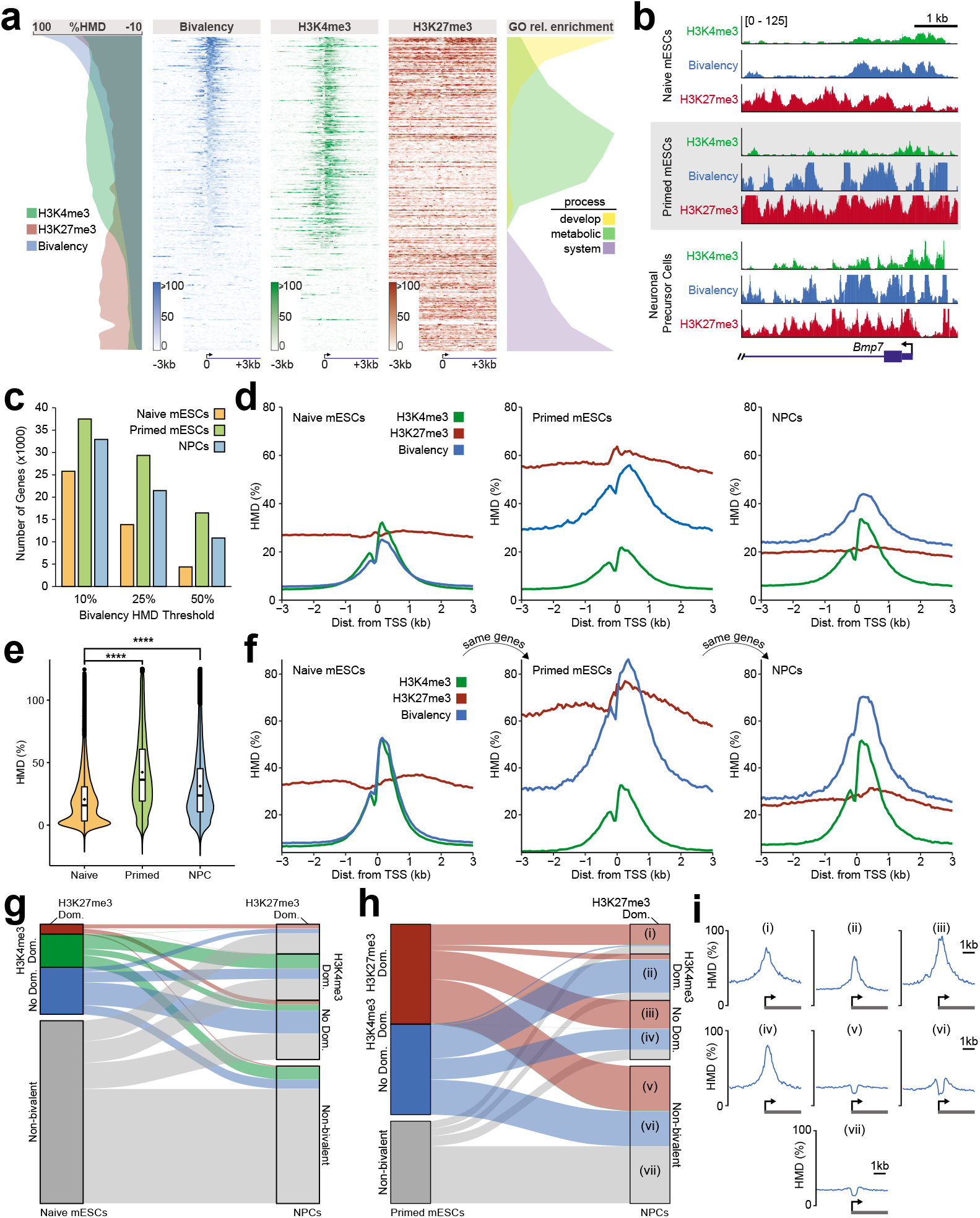
Bivalency is widespread and does not resolve over differentiation. **(a)** Bivalency, H3K4me3, and H3K27me3 at all Refseq promoters in naïve mESCs, with relative enrichment of GO terms. Genes are rank ordered by bivalency HMD at promoter, defined as the region from 0 to +400 bp relative to the TSS. **(b)** Representative locus view of H3K4me3, H3K27me3, and bivalency at promoters in naïve mESCs (top), primed mESCs (centre), and NPCs (bottom), presented on the same scale of 0-125% HMD. **(c)** Number of promoters with bivalency HMDs above the given thresholds in each cell type out of a total of 42,622 Refseq promoters. **(d)** Metaprofiles of H3K4me3, H3K27me3, and bivalency at all promoters in naïve mESCs, primed mESCs, and NPCs. Heatmaps for primed mESCs and NPCs are presented in Extended Data Fig. 3b. **(e)** Distribution of bivalency HMDs at all Refseq promoters in three cell states, zoomed to below 125% HMD. Overall, 99.5% of naïve promoters, 87.3% of primed promoters, and 91.6% of NPC promoters have an HMD below 100%. Full plot in Extended Data Fig. 3a. **(f)** Metaprofiles of H3K4me3, H3K27me3, and bivalency at promoters identified as bivalent in naïve mESCs (25% HMD threshold), tracked from naïve mESCs to primed mESCs to NPCs. Heatmaps for bivalency are presented in Extended Data Fig. 3f. **(g-h)** Alluvial plots of dominance and bivalency of genes from (g) naïve mESCs to NPCs or (h) primed mESCs to NPCs. Bivalency [>25% HMD] can be subcategorized into dominance classes by independent ICeChIP for the constituent marks, with H3K27me3 in excess (H3K27me3/H3K4me3 > *e^1^*), H3K4me3 dominant (H3K27me3/H3K4me3 < *e^−1^*), or intermediate ratios (no dominance). **(i)** Bivalency metaprofiles for gene subsets indicated in panel (h) from −3kb to +3kb relative to the TSS. **** *p* < 2.2 x 10^−16^.

Even more striking were the changes in bivalency across this differentiation scheme. Previous studies suggested that bivalency largely disappears upon differentiation to NPCs^16,18–20^. However, we found the opposite; promoter bivalency *increases* upon differentiation (Fig. 2d-e; Extended Data Fig. 3a-b), with thousands more genes meeting bivalency HMD thresholds relative to naïve mESCs (Fig. 2c). Similarly, we find that bivalent domains do not resolve upon differentiation; tracking bivalent genes from naïve mESCs through differentiation, we observe that bivalency is higher at these same promoters in primed mESCs and NPCs (Fig. 2f; Extended Data Fig. 3c-f). As previously reported, primed mESCs have the most bivalency, likely related to the high level of promoter H3K27me3 in this state^57^ (Fig. 2d). Accordingly, there are 27% fewer bivalent genes in NPCs than in primed mESCs (Fig. 2e). However, this decrease is nowhere near the previously reported decrease of 92%^18^, and bivalent genes from primed mESCs remain highly bivalent in NPCs (Extended Data Fig. 3g-h). Collectively, these data suggest that bivalency is far more widespread in this system than previously appreciated and remains elevated through differentiation, rather than resolving to one of the two monovalent states.

To investigate this discrepancy with the literature, we compared promoters identified as bivalent by other studies^18,20,35^ to ours. The previously identified genes had 50-100% more H3K27me3 than do most bivalent genes in our set (Extended Data Fig. 4a-b), suggesting that the previous studies undersampled H3K27me3 and thus could only identify regions with high H3K27me3 as bivalent. Accordingly, H3K27me3 dominant bivalent genes had the greatest proportional overlap with these canonical bivalent loci compared to other dominance classes (i.e. whether the bivalent genes have excess H3K27me3, excess H3K4me3, or roughly equal levels as measured by independent ICeChIP experiments for these two marks; Extended Data Fig. 4c). The common practice of measuring bivalency as regions of overlapping H3K4me3 and H3K27me3 is also problematic, even with calibrated data^46^; many promoters with high H3K4me3 and H3K27me3 bear less than 25% bivalency (Extended Data Fig. 4d). Notably, even for the previously identified bivalent genes, bivalency still increases relative to naïve mESCs upon differentiation. And in our datasets, this holds true across modification dominance classes – even the H3K27me3 dominant bivalent genes, which most closely resemble the canonically bivalent loci (Extended Data Figs. 4-5). To the extent that any bivalency class resolves from naïve mESCs to NPCs, the largest set of genes is from the H3K4me3 dominant bivalent genes (*p* = 1.78 x 10^−133^; Fig. 2g), despite its minimal overlap with the canonical bivalent loci (Extended Data Fig. 4c).

Having found that bivalency is unexpectedly common and persistent in early differentiation, we investigated the enzyme complexes that could potentially account for this ubiquity. Previous work suggested that H3K27me3 and H3K4me3 each inhibit deposition of the other^49,56,59,60^, particularly when symmetric (Supplementary Note 4), raising questions as to whether the pervasive bivalency we observe is plausible. To address this concern, we performed histone methyltransferase (HMTase) assays with Set1B and the full panel MLL-family core complexes (MLL1, MLL2, MLL3, MLL4), which collectively account for the bulk of H3K4 methylation in humans^61^. We find that these complexes all tolerate a wide spectrum of H3K27me3-decorated nucleosomes (Extended Data Fig. 6), indicating that the formation of bivalent nucleosomes is not precluded by allosteric modulation of H3K4me3 installation by core factors. Although it has been suggested that Set1a^62^, Mll2^63^, Ezh1^64^, and Ezh2^65^ are all important for establishing bivalency, only Mll2 appears to be sensitive for identifying bivalent promoters in naïve mESCs, with none showing high specificity for the same (Extended Data Fig. 7). Together, these data support the proposed specialized role for Mll2 in bivalency^63^, indicate a pleiotropic role for PRC2 beyond its role in establishing bivalency, and provide plausible enzymatic avenues to the prevalent bivalency we observe by reICeChIP.

## Bivalency, gene expression, and ontology

A key pillar of the bivalency hypothesis is that bivalent promoters are associated with transcriptionally repressed genes poised to either be activated or terminally silenced upon differentiation^16,18,21,22^. However, bivalency is not solely found at genes with low expression in any of our measurements (Fig. 3a; Extended Data Fig. 8a-b). Rather, bivalent genes had higher average expression than did non-bivalent genes or the set of all genes, and these genes display modestly higher average expression through differentiation (Fig. 3a), with bivalency remaining similar across most gene expression deciles (Extended Data Fig. 8c). Bivalency associated similarly with bulk gene expression (Fig. 3b) and the proportion of cells expressing the associated transcripts in single cell RNA-seq (Fig. 3c), suggesting that the association of bivalency with higher-expressed genes is not solely driven by intercellular heterogeneity. Consistent with previous observations^18^, bivalency was higher at promoters with high CpG content (Extended Data Fig. 8d) and associated with lower DNA methylation compared to non-bivalent genes (Extended Data Fig. 8e, also holds for each dominance class). These data all suggest that bivalent genes are more highly expressed than non-bivalent genes as a whole, and this latter class is seemingly more subject to regulation by DNA methylation.

**Fig. 3.**
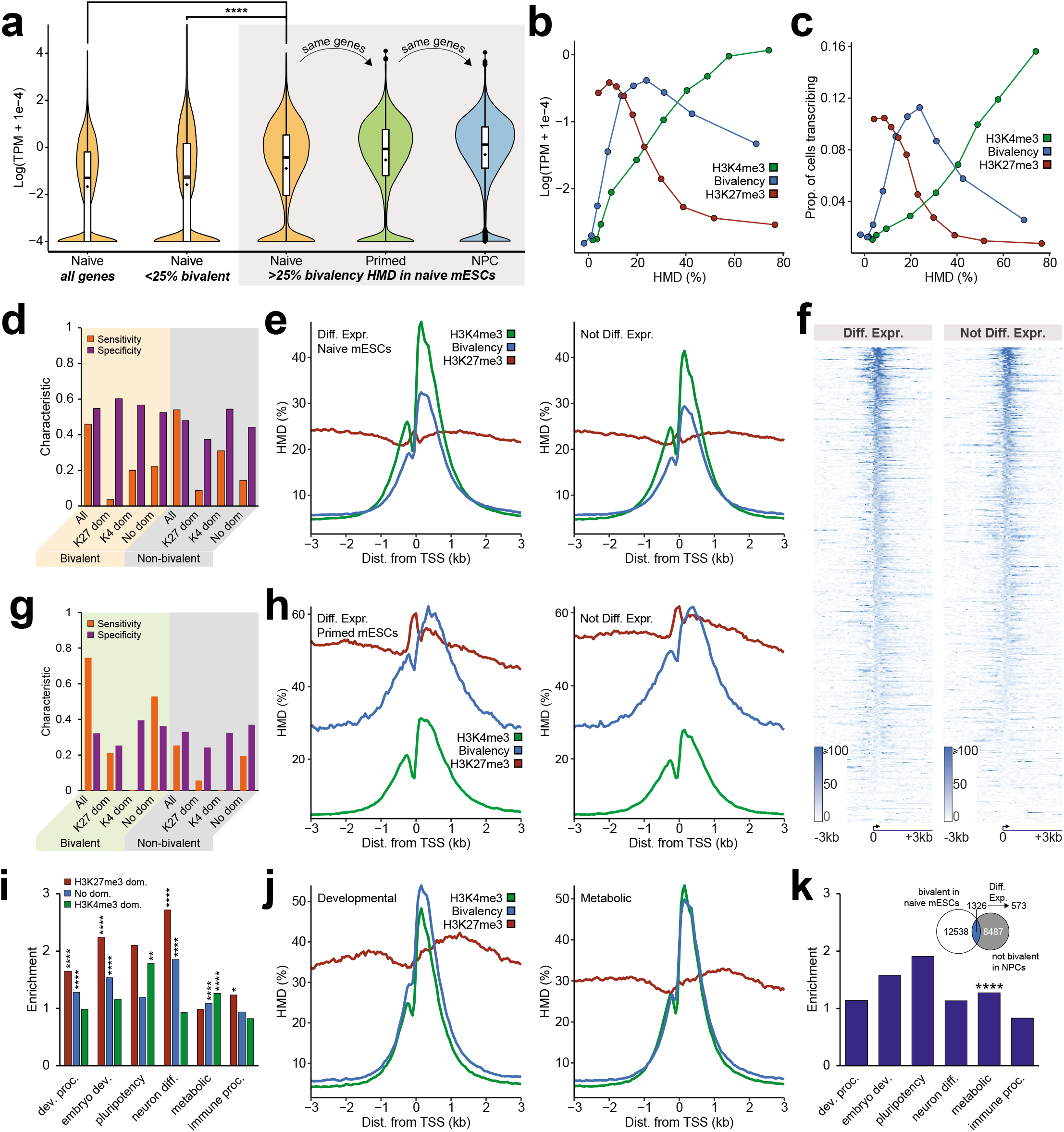
Bivalency is neither sensitive nor specific for poised nor developmental genes. **(a)** Violin plots of gene expression^79^ for all genes in naïve mESCs, non-bivalent genes (<25% HMD) in naïve mESCs, and bivalent genes (>25% HMD) tracked from naïve mESCs to the same genes in the indicated lineages. Significance computed by Welch’s two-tailed *t*-test. **(b)** Gene expression vs. HMD for H3K4me3, H3K27me3, and bivalency (genes are binned into HMD deciles). **(c)** Proportion of actively transcribing cells by single-cell RNA-seq^80^ vs. HMD for H3K4me3, H3K27me3, and bivalency (genes are binned into HMD deciles). **(d)** Sensitivity and specificity (Supplementary Note 5) of bivalent and non-bivalent genes in naïve mESCs identifying differentially expressed genes (DEGs) from the naïve state to the NPC state. **(e)** Metaprofiles of H3K4me3, H3K27me3, and bivalency and **(f)** heatmaps of bivalency in naïve mESCs at DEGs and non-DEGs relative to NPCs. **(g)** Sensitivity and specificity of bivalent and non-bivalent genes in primed mESCs identifying DEGs from the primed state to the NPC state. **(h)** Metaprofiles of H3K4me3, H3K27me3, and bivalency in primed mESCs at DEGs and non-DEGs. **(i)** Gene ontology term enrichment of H3K27me3-dominant bivalent genes, H3K4me3-dominant bivalent genes, or bivalent genes with no clear dominance (q-value two-tailed Fisher hypergeometric test). **(j)** Metaprofiles of H3K4me3, H3K27me3, and bivalency in naïve mESCs at developmental and metabolic genes. **(k)** Gene ontology term enrichment of genes following the classic bivalency model: DEGs that lose bivalency from naïve mESCs (>25% HMD) to NPCs (<10% HMD). Significance computed by two-tailed Fisher hypergeometric test. * *q* < 0.05. ** *q* < 0.01. **** *p* or *q* < 2.2 x 10^−16^.

Another pillar of the bivalency model is that bivalent genes are poised to be differentially regulated through differentiation. To test this, we computed the sensitivity and specificity of different bivalency and non-bivalency classes for differentially expressed genes (DEGs; Supplementary Note 5). Counter to the bivalency hypothesis and previous results^16,18,21^, we found that bivalency was a very poor marker of DEGs; from naïve mESCs to NPCs, bivalency was roughly as sensitive and specific for identifying DEGs as was a *lack* of bivalency (Fig. 3d). Though H3K27me3-dominant bivalent genes showed an increase in average gene expression (Extended Data Fig. 8f-g), this class still only had 60% specificity for identifying DEGs, with very low sensitivity (Fig. 3d). Promoters of DEGs and non-DEGs from naïve mESCs to NPCs have highly similar histone modification metaprofiles in naïve mESCs (Fig. 3e-f) and across differentiation (Extended Data Fig. 8h-k). Comparison of primed mESCs to NPCs displayed similar trends (Fig. 3g-h); though sensitivity was higher because most genes are bivalent in primed mESCs, the specificity remained similar between bivalent and non-bivalent genes. Interestingly, whether genes were upregulated, downregulated, or non-DEGs, bivalency still increased over differentiation (Extended Data Fig. 8h-k). Collectively, these analyses show that bivalency is neither sensitively nor specifically associated with poised DEGs in this system.

We next examined whether bivalency is primarily associated with developmental genes, a central tenet of the original model^16,18^. The first ICeChIP study indirectly hinted that there may be at least two classes of bivalent promoters: an H3K27me3 dominant class associated with developmental genes, and an H3K4me3 dominant class enriched for metabolic genes^46^. Direct measurements of bivalency herein unambiguously demonstrate this phenomenon more broadly (Fig. 2a, 3i). Overall, bivalent genes are enriched for a broad range of ontology terms, including developmental, metabolic, and immune system process genes (Fig. 3i-j), with nearly identical bivalency profiles in naïve mESCs (Fig. 3i; Extended Data Fig. 9a). These classes all not only retained, but increased bivalency into NPCs – even immune system process genes, despite being seemingly unrelated to neuronal development. We only found 543 genes that *did* obey the classic bivalency model (Fig. 3k), representing less than 5% of the bivalent genes from naïve mESCs, with little difference in bivalency between upregulated and downregulated genes (Extended Data Fig. 9b). Interestingly, these genes were most significantly enriched for metabolic rather than developmental genes (Fig. 3k). Taken together, these data suggest that bivalency is neither primarily nor specifically associated with developmental genes in this system.

## Predicting DEGs with histone PTMs

The premise of the bivalency hypothesis is that the coexistence of H3K4me3 and H3K27me3 synergistically provides additional predictive information about the associated genes upon differentiation beyond that provided by H3K4me3 and H3K27me3 alone. With quantitative measurements of these modifications, this hypothesis can be tested by modelling. We first determined which individual parameters best identified DEGs by measuring the area under the curve (AUC) of receiver operator characteristic (ROC) curves of parameter thresholds. Of the individual histone modifications, H3K4me3 levels were best for identifying DEGs, with the highest AUC of the ROC (Fig. 4a; Extended Data Fig. 10a). Bivalency was less predictive of DEGs than were either the log ratio of H3K27me3 and H3K4me3 or DNA methylation (Fig. 4a; Extended Data Fig. 10a). And in primed mESCs, far from being predictive of poised genes, bivalency was *inversely* associated with DEGs upon differentiation to NPCs (Extended Data Fig. 10a).

**Fig. 4.**
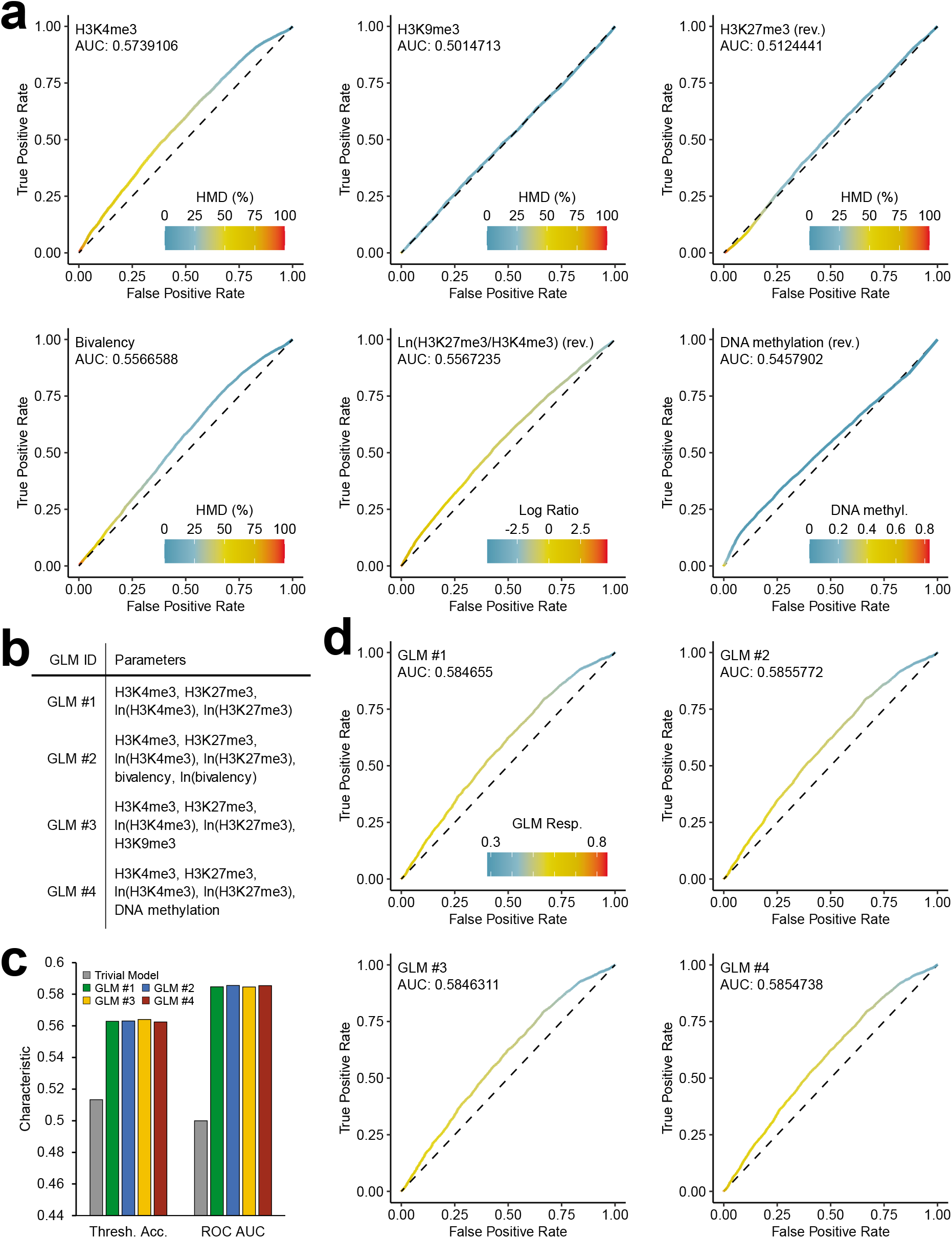
Bivalency does not provide appreciably more information than H3K4me3 and H3K27me3 alone for DEG prediciton. **(a)** Receiver operator characteristic (ROC) curves for identifying DEGs from naïve mESCs to NPCs by H3K4me3, H3K9me3, H3K27me3, bivalency, ln(H3K27me3/H3K4me3), or DNA methylation in naïve mESCs. For each point, parameter value threshold used to compute true positive rate (TPR) and false positive rate (FPR) is indicated by the colour. Traits with thresholds identifying non-DEGs rather than DEGs are marked with “rev.” **(b)** Legend for generalized linear models (GLMs) in panels c-d. **(c)** Accuracy of trivial model and GLMs by threshold accuracy (gene identified as DEG if logistic regression > 0.5; left) and by ROC area under curve (right). **(d)** ROC curves for identifying DEGs from naïve mESCs to NPCs by different GLMs. For each point, logistic regression threshold value used to compute TPR and FPR is indicated by the colour.

If bivalency provides additional information over H3K4me3 and H3K27me3, then a model without bivalency will be markedly less explanatory than a model with bivalency. To test this, we conducted logistic regressions with linear models to identify parameters most important for identifying DEGs. Bayes Information Criterion analyses preliminarily hinted that bivalency provided minimal information to this end (Extended Data Fig. 10b; Supplementary Note 6). To more definitively identify whether bivalency provides meaningful predictive information, we conducted hold-out cross-validation on models with H3K4me3, H3K27me3 and either nothing else, bivalency, H3K9me3, or DNA methylation (Fig. 4b; Extended Data Fig. 10c; Supplementary Note 6). Parameters other than H3K4me3 and H3K27me3 barely improved model accuracy by two separate metrics (Fig. 4c-d; Extended Data Fig. 10d-e; Supplementary Note 4), suggesting that those parameters provide virtually no additional information content to identify DEGs. These data suggest that, in this developmental system, there is little evidence that bivalency has emergent properties in identifying poised genes beyond the combined independent properties of H3K4me3 and H3K27me3.

## Discussion

The bivalency hypothesis is one of the more influential ideas in epigenetics and molecular developmental biology. Persistent interest over the years coupled with widespread deployment and acceptance of sub-optimal bivalency measurement methods has ossified the hypothesis into dogma that extends well beyond any of the experimental data that informed it.

However, this coalescence has not been reached based on functional assays. Indeed, to the extent that functional validation of the bivalency model has been attempted, it has primarily been through deletion of enzymes with pleiotropic effects and functions throughout the genome beyond installation of bivalency^35,63,66,67^. Overwhelmingly, the prevailing views on the role of bivalency are derived from ChIP experiments. However, ChIP protocols^68^ and antibodies^47,69–72^ are often highly susceptible to off-target pulldown, and uncalibrated ChIP without exogenous normalization can distort signal and the ability to compare experiments^46,47,73^, leading to spurious conclusions^47^. From the quantitative and specific measurements we made with reICeChIP, we fear that this has been the case with the bivalency hypothesis, at least as far as these analyses in early mESC differentiation permit.

It has been held that bivalency is present at a small, restricted set of promoters early in development; we find that bivalency is widespread, with many thousands of promoters displaying high bivalency levels. It has been held that bivalency primarily exists early in development and resolves upon differentiation; we find that bivalency persists at least through the NPC stage and *increases* over baseline in that span. It has been held that bivalency demarcates poised, developmental genes associated with lineage commitment; we find that bivalency is neither sensitively nor specifically associated with developmental nor differentially expressed genes – and, at worst, may be *inversely* associated with the latter. Moreover, bivalent genes are predominantly not poised in an off state, but are more highly expressed than those that are not bivalent. All told, we find little evidence that bivalency provides more information in predicting poised gene status than do H3K4me3 and H3K27me3 in an independently additive manner in this system, raising questions as to whether it represents any more than a coincidental overlap of the aforementioned two marks.

Our study is not without caveats. First, we are only able to comment meaningfully on the differentiation paradigm presented here; we cannot definitively infer that these results will hold for the other developmental or clinical contexts. Although the original studies on bivalency indicated that bivalency almost entirely disappeared by the NPC stage^16,18^, this stage is not terminally differentiated, so it is possible that bivalency could resolve in later stages of differentiation. Future studies will be needed to address this possibility in other developmental contexts. Second, though the extant evidence suggests that only trans-bivalency is present at meaningful levels, our method cannot selectively distinguish between *cis*-, *trans*-, and intermediate bivalency conformations (Supplementary Note 1).

The reICeChIP method is not inherently restricted to the study of H3K4me3/H3K27me3 bivalency. With cleavable recombinant affinity reagents targeting other histone modifications^53,74^ it could be used to quantify other combinatorial modification patterns^75–78^ or modification symmetry.

Without serious changes to the standards of ChIP, the limitations of conventional ChIP-seq will continue to pose an existential challenge to the field. Indeed, the divergence between our observations of bivalency and those in the literature can be attributed to the historical lack of tools needed to make quantitative and specific measurements; in that context, the experimental designs and interpretations of the past were reasonable. Fortunately, such tools now exist. And as we have shown in this work, these methods offer a chance for the field to critically evaluate its orthodox models and pave the way for new insights on the chromatin determinants of cell identity and the regulation of development.

## Acknowledgements

We would like to thank Peter Faber, Hannah Whitehurst, and Mikayla Marchuk in the University of Chicago Functional Genomics Facility for Illumina sequencing. We would also like to thank EpiCypher, Inc. for providing some of the histone octamers for this study. A.T.G. was supported by the Harper Dissertation Prize and the Dean’s International Student Fellowship of the University of Chicago. R.N.S. was supported by the National Institutes of Health under award number T32-HD007009-45 to the University of Chicago. J. E. was supported by the National Institutes of Health under award number T32-GM007197 and R25-GM109439 to the University of Chicago. This study was supported by the National Institutes of Health, under award numbers R01-GM115945 to A.J.R. and R01-DA036887 to S.K.; and the American Cancer Society, under award number 130230-RSG-16-248-01-DMC to A.J.R.

## Author Contributions

A.T.G. and A.J.R. conceived of the study. Z.C. made *cis*-bivalent H3K4me3-H3K27me3 histones. C.C.L. and B.F. constructed the *trans*-bivalent H3K4me3-H3K27me3 histone dimers. T.H. and S.K. provided 304M3B Fab and assisted in design of 304M3B-1HRV3C Fab. P.W.L. provided recombinant PRC2 for supporting experiments. A.T.G. developed reICeChIP, expressed and purified 304M3B-1HRV3C Fab, and conducted methyltransferase assays with input and oversight from A.J.R. A.T.G. cultured naïve mESCs; J.E. cultured and differentiated naïve mESCs to primed mESCs and NPCs. A.T.G. conducted reICeChIP-seq on naïve mESCs; R.N.S. conducted reICeChIP-seq on primed mESCs and NPCs. A.T.G. and R.N.S. conducted bioinformatic analysis on naïve mESCs; J.E. generated representative genome browser views. R.N.S. conducted all other bioinformatic analyses with oversight and input from A.T.G., J.E., and A.J.R. R.N.S., A.T.G., J.E., and A.J.R. wrote the manuscript with input from the other authors.

## Additional Information

**Supplementary Information** is available for this paper.

**Correspondence and requests for materials** should be addressed to A.J.R.

**Reprints and permissions information** is available at http://www.nature.com/reprints.

## Extended Data Figure Captions

**Extended Data Fig. 1.**
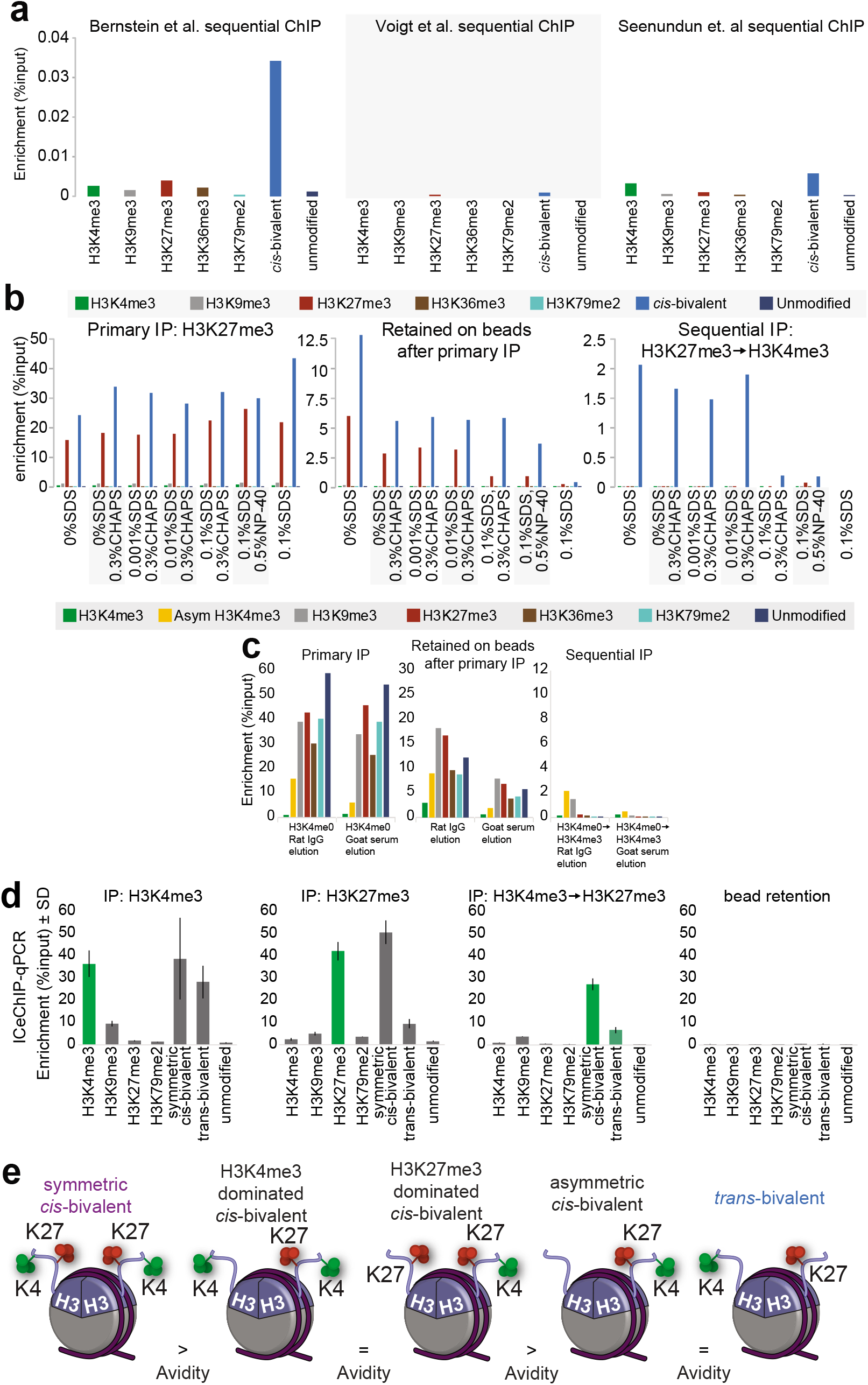
Evaluation of sequential ChIP methods. **(a)** Enrichment of on- and off-target nucleosome standards under sequential ChIP protocols developed by Bernstein et al.^16^, Voigt et al.^49^, and Seenundun et al.^29^. **(b-c)** Enrichment at different sequential ICeChIP steps with (b) chemical denaturant elution and (c) immunoglobulin and serum elution. **(d)** Enrichment of different nucleosome standards with ICeChIP-qPCR performed against H3K4me3, H3K27me3, and bivalency, with beads showing very little retention of chromatin (n=3 technical replicates). Error bars represent standard deviation. **(e)** Different configurations of bivalency on a single nucleosome. Of these, only trans-bivalency has been identified by mass spectrometry^49,81^.

**Extended Data Fig. 2.**
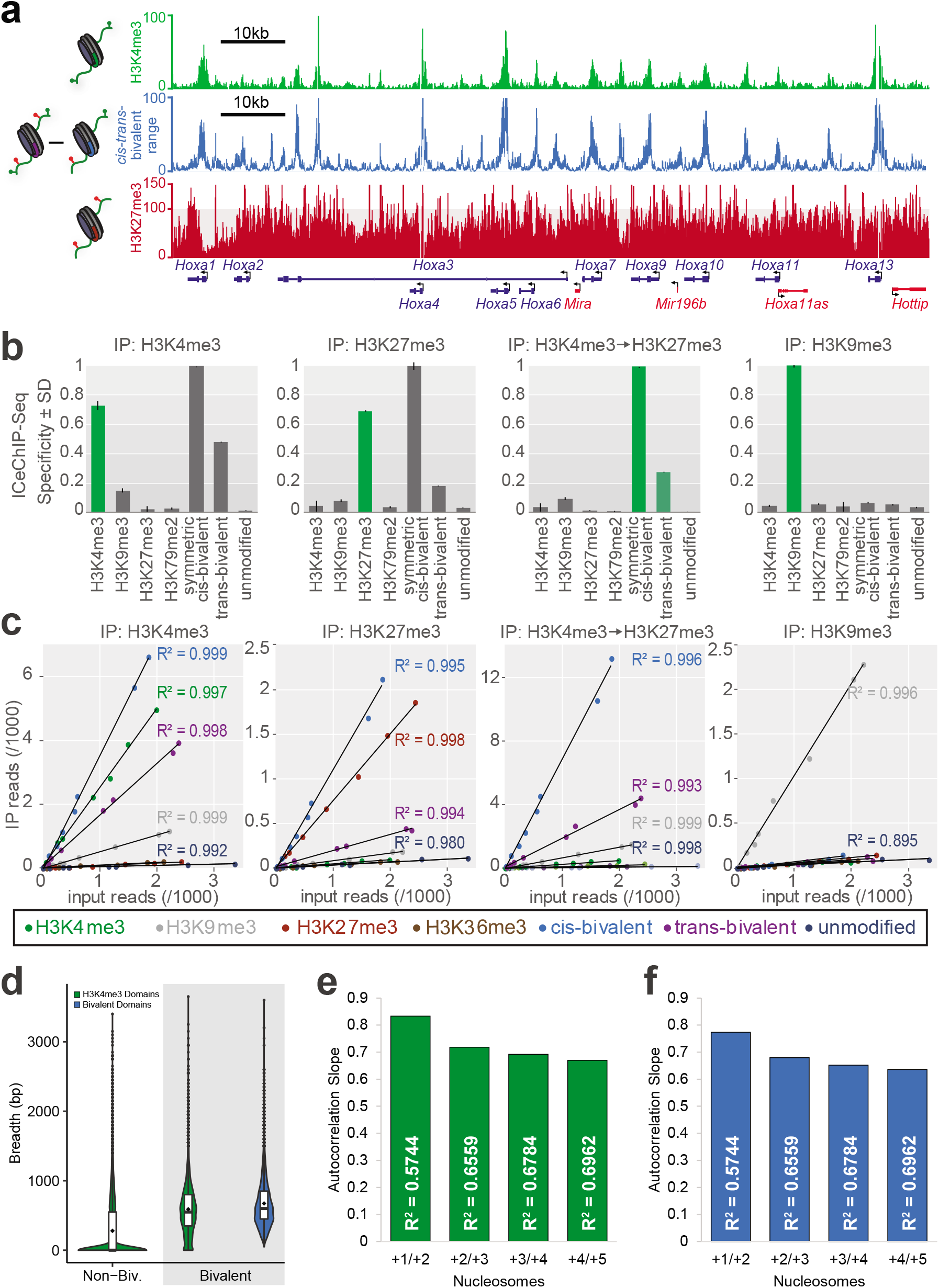
Evaluation of reICeChIP specificity and standards. **(a)** Representative genome browser view of H3K4me3, H3K27me3, and bivalency, shown as a range of possible values by normalization to trans-bivalent (upper limit) or cis-bivalent (lower limit) nucleosome standards. **(b)** Relative pulldown of different nucleosome standards in ICeChIP-seq, normalized to the most-enriched standard. **(c)** Scatterplots of reads from DNA barcodes applied to nucleosome standards in ICeChIP-seq. **(d)** Violin plots of peak breadth (consecutive segment of 50bp windows overlapping promoter with >25% HMD) for H3K4me3 (green) and bivalency (blue) at non-bivalent and bivalent genes (>25% HMD) in naïve mESCs. **(e-f)** Autocorrelation of (e) H3K4me3 and (f) bivalency HMDs between nucleosomes in naïve mESCs. Nucleosomes are defined as sequential 200bp windows from the TSS.

**Extended Data Fig. 3.**
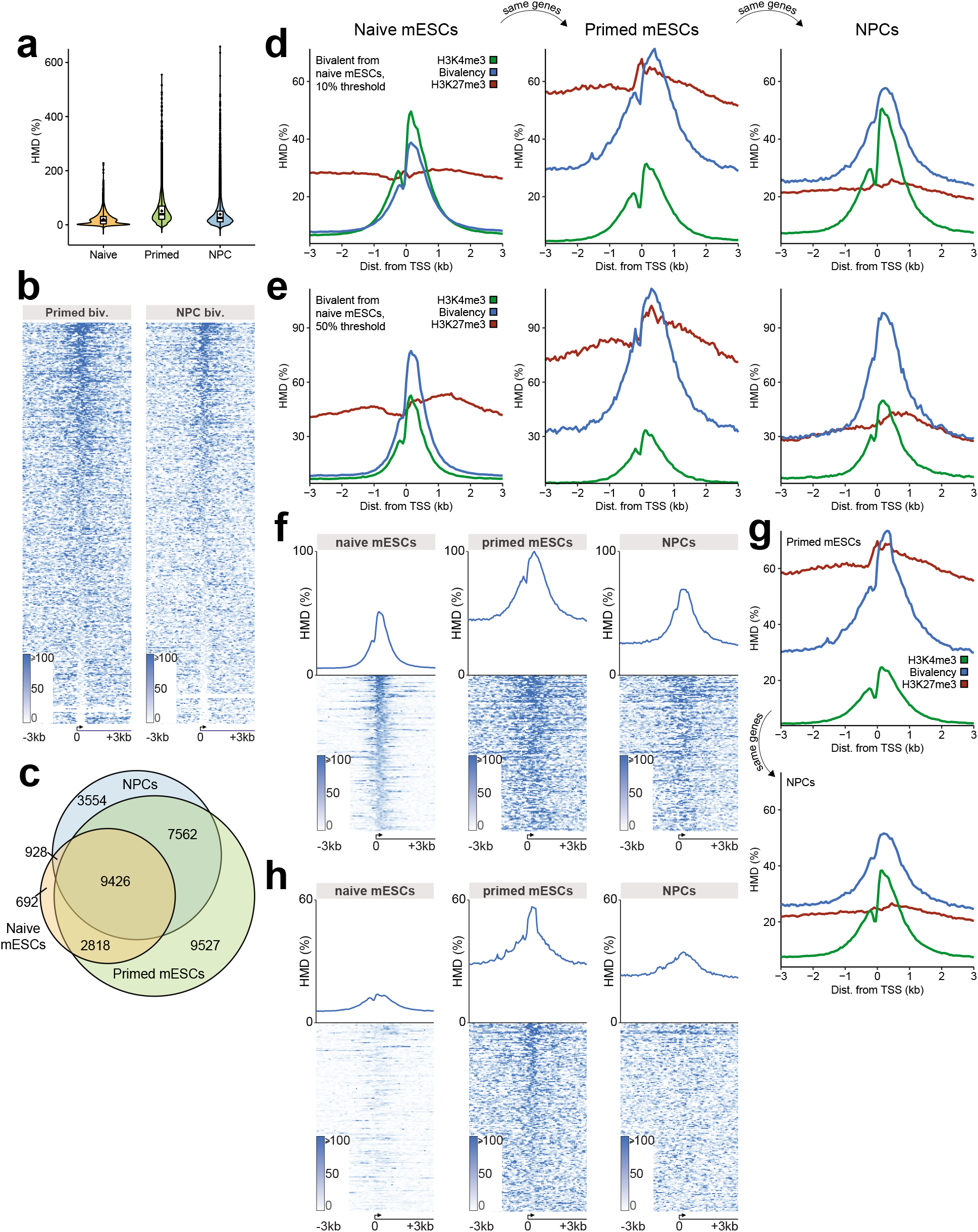
Tracking bivalent genes through differentiation. **(a)** Distribution of bivalency HMDs at all Refseq promoters in three cell states. **(b)** Heatmaps of bivalency at all Refseq promoters in primed mESCs and NPCs. Genes are ordered by bivalency HMD at the promoter. **(c)** Venn diagram showing overlap of bivalent genes (25% HMD threshold) in naïve mESCs, primed mESCs, and NPCs. **(d-e)** Metaprofiles of H3K4me3, H3K27me3, and bivalency for bivalent genes in naïve mESCs with a (d) 10% or (e) 50% HMD threshold. **(f)** Heatmaps and metaprofiles of bivalent genes from naïve mESCs. **(g)** Metaprofiles of H3K4me3, H3K27me3, and bivalency at genes tracked from primed mESCs to NPCs for bivalent genes in primed mESCs (>25% HMD). **(h)** Heatmaps and metaprofiles of bivalent genes in primed mESCs that are not bivalent in naïve mESCs.

**Extended Data Fig. 4.**
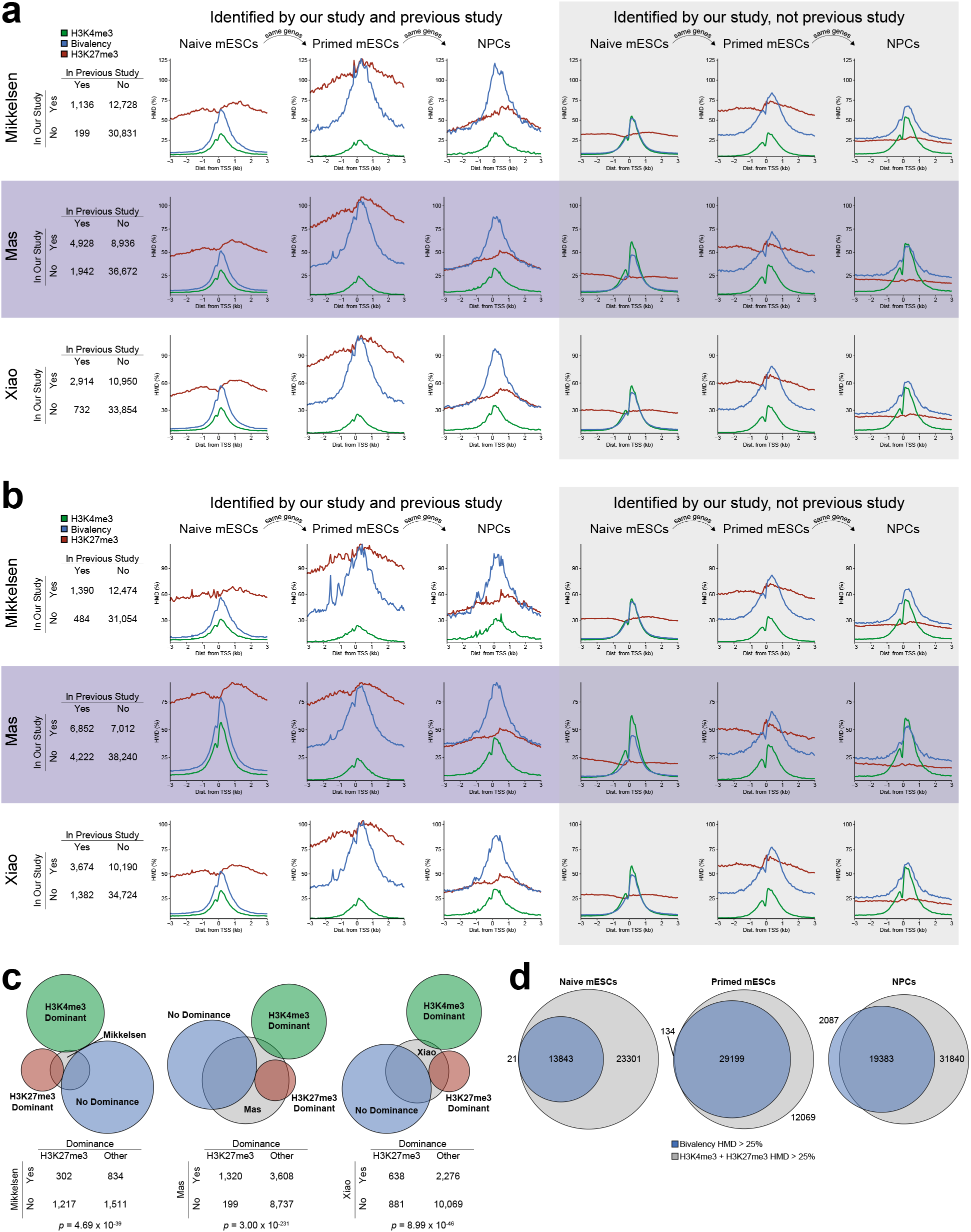
Comparing our bivalent genes to other studies. **(a-b)** Contingency tables and metaprofiles for genes that are identified as >25% bivalent in our study and by Mikkelsen et al.^18^, Mas et al.^35^, and Xiao et al.^58^, wherein: (a) gene is identified as bivalent in the external study if overlapping H3K4me3 and H3K27me3 peaks overlap the 0 to +400bp region of a gene relative to the TSS, or (b) gene is identified as bivalent in the external study if overlapping H3K4me3 and H3K27me3 peaks overlap the region from 2.5kb upstream of the TSS to the end of the gene^22^. **(c)** Overlap of bivalent genes from external datasets (as defined in part a) with each of our bivalent gene dominance classes in naïve mESCs. Significance computed by two-tailed Fisher hypergeometric test. **(d)** Overlap of genes with bivalency HMD > 25% and with H3K4me3 + H3K27me3 HMD > 25% in all three cell states.

**Extended Data Fig. 5.**
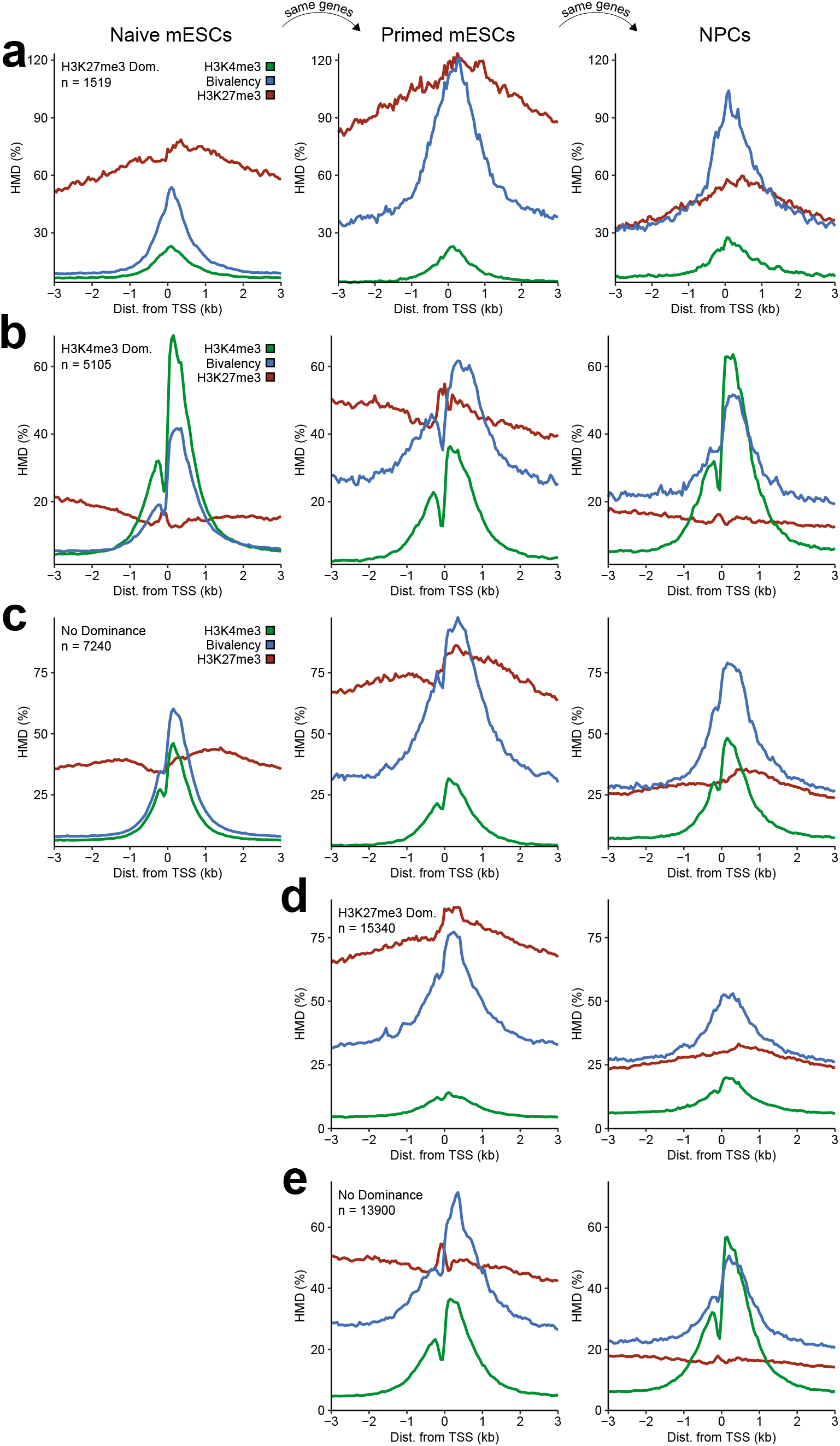
Bivalency changes across differentiation by modification dominance class. **(a-c)** Metaprofiles of H3K4me3, H3K27me3, and bivalency for bivalent genes (>25% HMD) in naïve mESCs that are (a) H3K27me3 dominant (H3K27me3/H3K4me3 > *e^1^*), (b) H3K4me3 dominant (H3K27me3/H3K4me3 < *e^−1^*), or (c) have no dominance in naïve mESCs, tracked through three cell states. **(d-e)** Metaprofiles of H3K4me3, H3K27me3, and bivalency for bivalent genes (>25% HMD) in primed mESCs that are for indicated dominance classes tracked from primed mESCs to NPCs.

**Extended Data Fig. 6.**
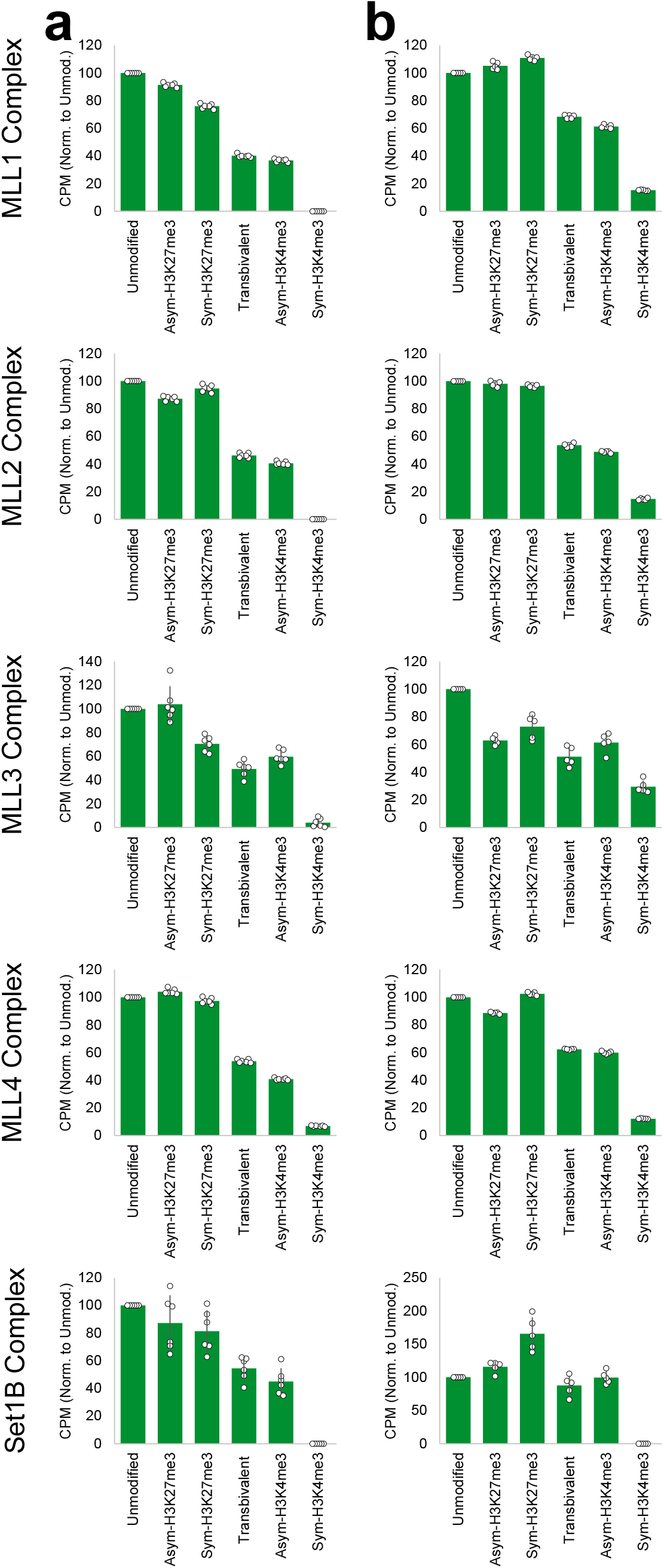
Methyltransferase assays identifying potential pathways for establishment of bivalency. **(a-b)** Methyltransferase assays for MLL1, MLL2, MLL3, MLL4, and Set1B core HMTase complexes using (a) 15 ng/uL (n=6) and (b) 20 ng/uL (n=5) semisynthetic nucleosomes as substrates for methylation. Endpoints were established at 180 min by kinetic evaluation to be sensitive to difference in activity for this panel. Signal is corrected for background and no nucleosome substrate activity. Error bars represent standard deviation.

**Extended Data Fig. 7.**
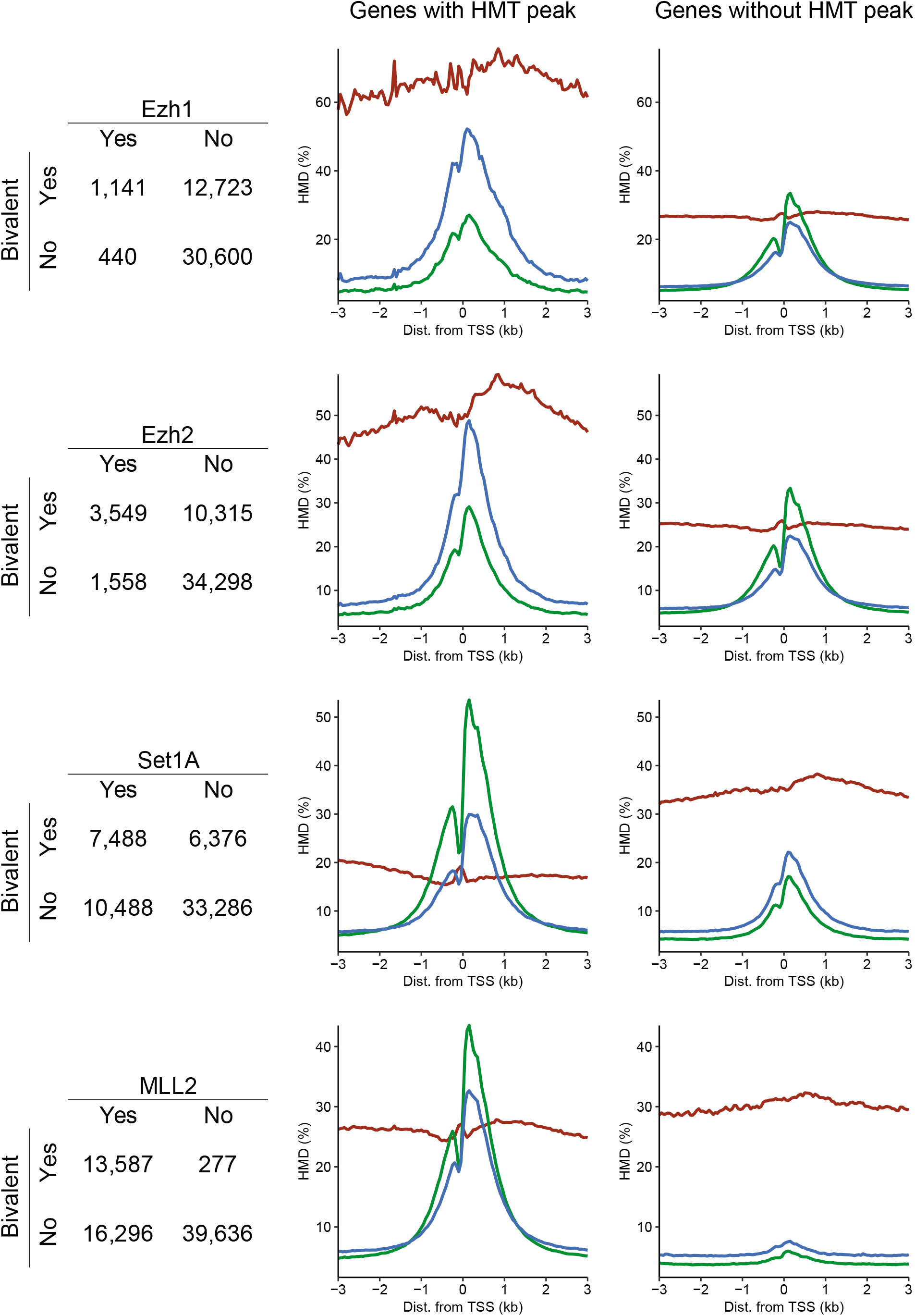
HMTase peaks and bivalency. Contingency tables and metagene profiles in naïve mESCs for genes with and without overlapping HMT peaks. Ezh1 and Ezh2 peaks were identified as Suz12 peaks lost upon Ezh1 or Ezh2 knockout^82^. Set1A peaks were identified by ChIP against Set1A^62^. Mll2 peaks were identified by ChIP against Mll2^45^.

**Extended Data Fig. 8.**
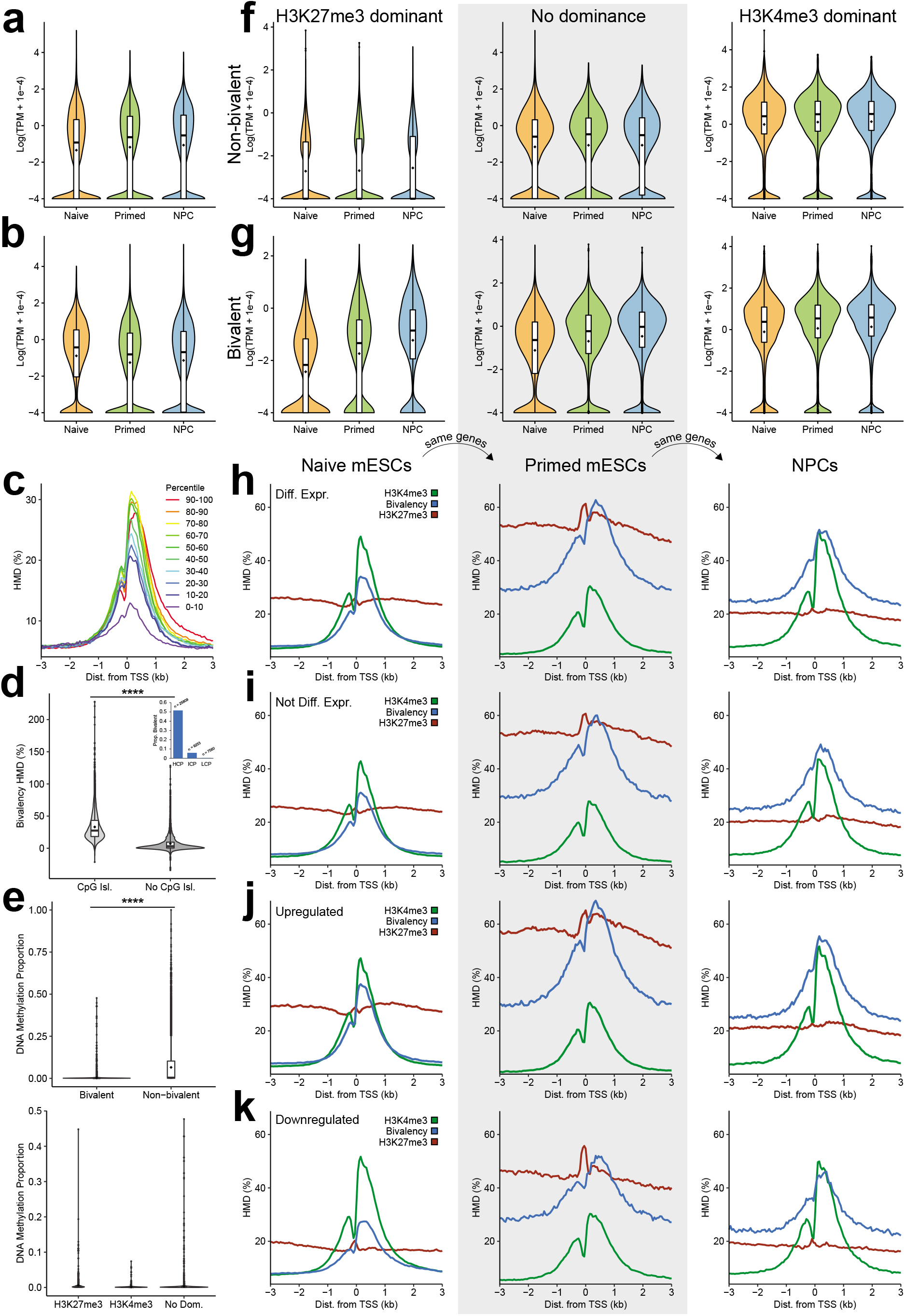
Bivalency and differential gene expression. **(a-b)** Violin plots of gene expression for (a) all genes and (b) bivalent (>25% HMD) genes in each cell state. **(c)** Bivalency metaprofiles in naïve mESCs at promoters binned by gene expression decile. **(d)** Violin plots of bivalency HMD in naïve mESCs at promoters with and without CpG islands. Inset shows proportion of genes that are bivalent in sets of genes classified by CpG content: high-CpG promoters (HCP), intermediate-CpG promoters (ICP), and low-CpG promoters (LCP), defined as previously described by Mikkelsen et al.^18^. Total number of genes in each class is provided as n. **(e)** Violin plots of DNA methylation at bivalent and non-bivalent genes (top), broken by dominance class for bivalent genes (bottom). **(f-g)** Violin plots of gene expression in (f) non-bivalent (<25% HMD) and (g) bivalent (>25% HMD) genes from naïve mESCs that are H3K27me3 dominant (H3K27me3/H3K4me3 > *e^1^*; left), have no clear dominance (centre), or are H3K4me3 dominant (H3K27me3/H3K4me3 < *e^−1^*; right). **(h-k)** Metaprofiles of H3K4me3, H3K27me3, and bivalency at genes tracked from naïve mESCs to primed mESCs to NPCs for (h) DEGs, (i) non-DEGs, (j) genes upregulated from naïve mESCs to NPCs, and (k) genes downregulated from naïve mESCs to NPCs. **** *p* <10^−16^ (Welch’s two-tailed t-test).

**Extended Data Fig. 9.**
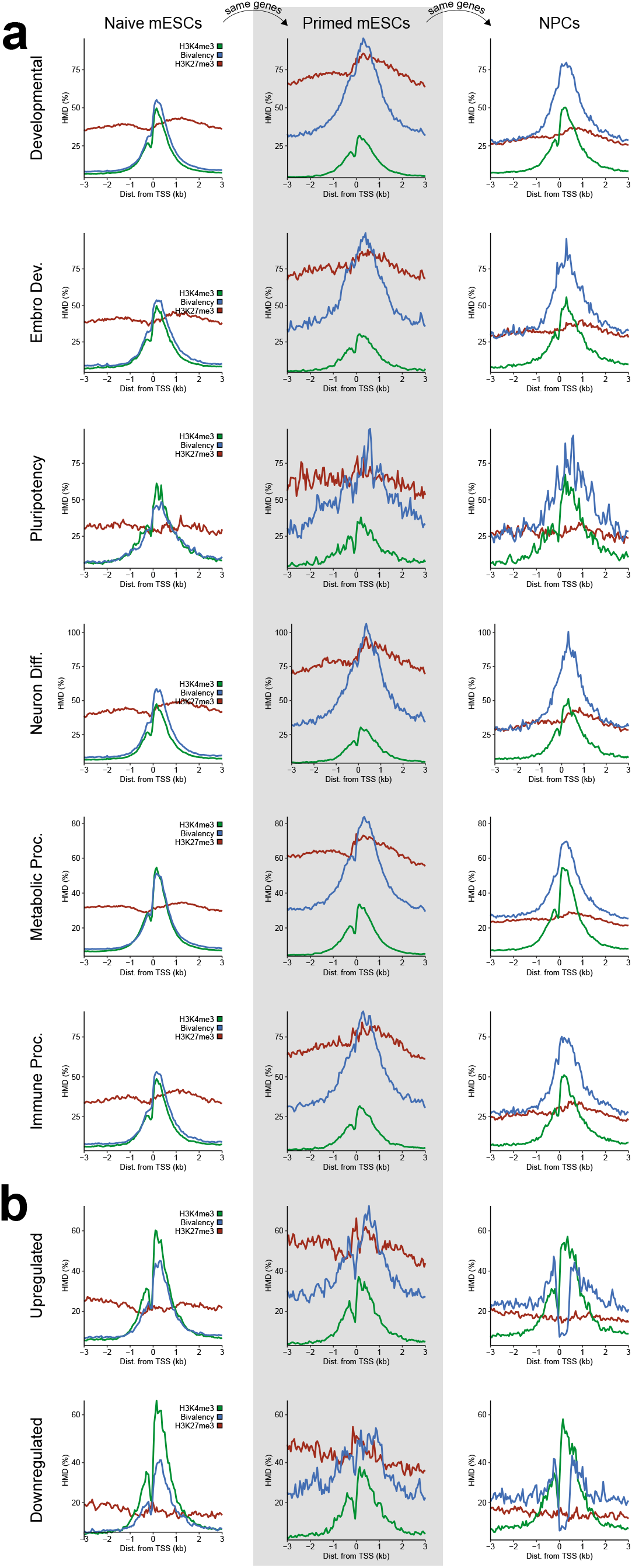
Bivalency at different classes of genes. **(a)** Metaprofiles of H3K4me3, H3K27me3, and bivalency at genes tracked from naïve mESCs to primed mESCs to NPCs for bivalent genes of indicated gene ontology terms. **(b)** Metaprofiles of H3K4me3, H3K27me3, and bivalency at genes tracked across differentiation for genes that lose bivalency at the promoters (0 to +400bp relative to TSS) from naïve mESCs (>25% HMD) to NPCs (<10% HMD) and are upregulated (top) or downregulated (bottom) over differentiation.

**Extended Data Fig. 10.**
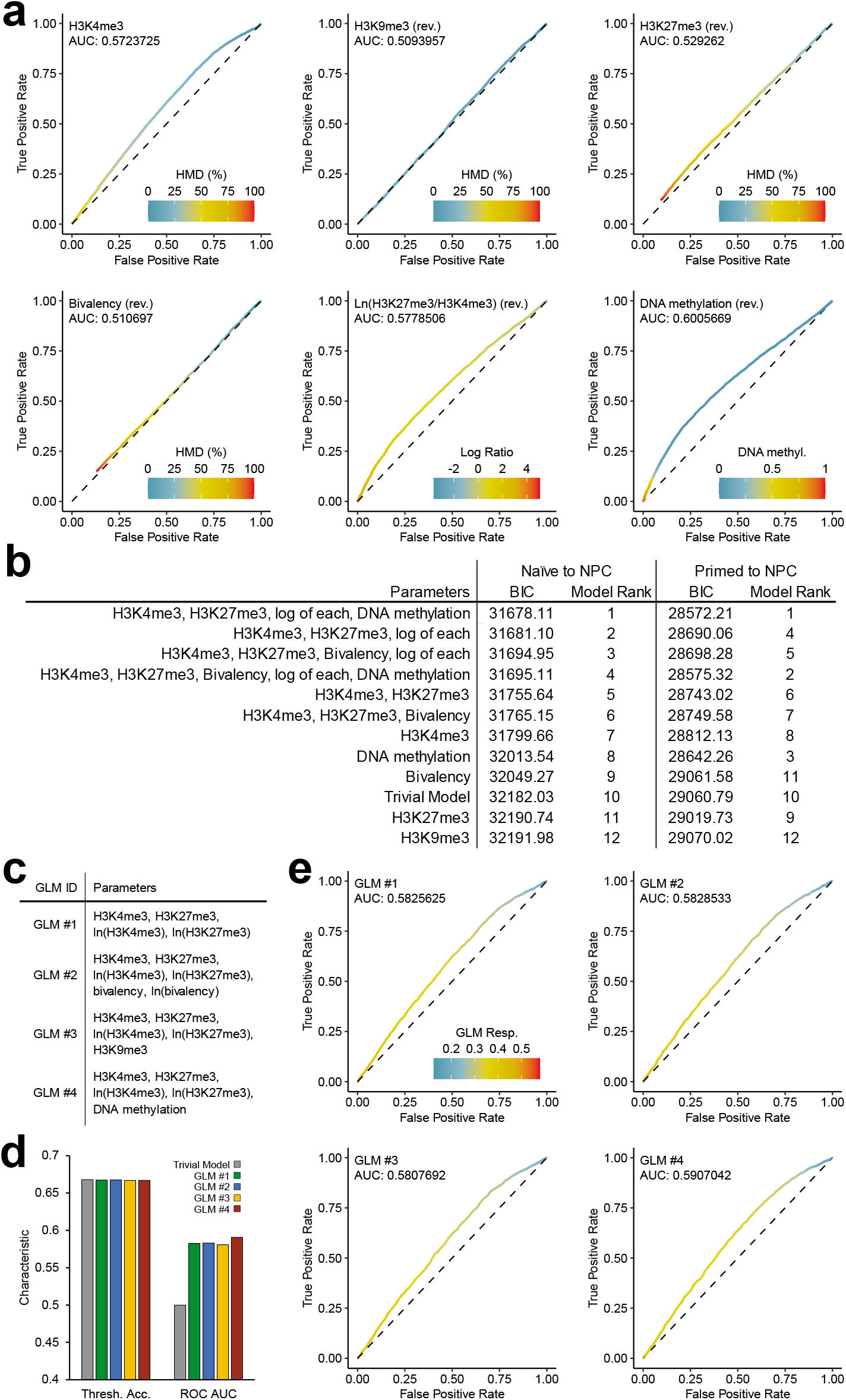
Modelling the additional information content provided by bivalency over H3K4me3 and H3K27me3 alone. **(a)** ROC curves for identifying DEGs from primed mESCs to NPCs by H3K4me3, H3K9me3, H3K27me3, bivalency, ln(H3K27me3/H3K4me3), or DNA methylation in primed mESCs. For each point, parameter value threshold used to compute true positive rate (TPR) and false positive rate (FPR) is indicated by the colour. Traits with thresholds identifying non-DEGs rather than DEGs are marked with “rev.” **(b)** Bayes Information Criterion (BIC) for logistic models identifying DEGs from naïve mESCs or primed mESCs to NPCs with different parameters. **(c)** Legend for generalized linear models (GLMs). **(d)** Accuracy of trivial model and GLMs by threshold accuracy (gene identified as DEG if logistic regression > 0.5; left) and by ROC area under curve (right). **(e)** ROC curves for identifying DEGs from primed mESCs to NPCs by different GLMs. For each point, logistic regression threshold value used to compute TPR and FPR is indicated by the colour.

## Supplementary Notes

### Supplementary Note 1

As each nucleosome has two H3 protomers, there are several different configurations of bivalency that a bivalent nucleosome can theoretically adopt, each with a different avidity for ChIP pulldown with immobilized antibody. At one extreme, with the highest avidity, is the symmetric *cis*-bivalency form, where both H3K4 and both H3K27 residues are trimethylated (Extended Data Fig. 1e). This nucleosome has the most epitopes for antibody binding and will thus have the highest avidity in pulldown reflected in apical pulldown efficiency (Extended Data Fig. 1d). At the other extreme, with the lowest avidity, is the *trans*-bivalency form, where single H3K4me3 and H3K27me3 marks decorate different histone tails (Extended Data Fig. 1e). This has the fewest epitopes for antibody binding and will thus have no avidity in pulldown.

This poses a theoretical challenge in normalization and calibration of a ChIP study; because we cannot separately measure *trans*-bivalency, symmetric *cis*-bivalency, nor any intermediate states, it is impossible for us to definitively state whether a given locus with a given HMD has relatively few nucleosomes that are symmetric *cis*-bivalent or whether it has relatively many nucleosomes that are trans-bivalently modified. To accommodate for this limitation, we include two different bivalent calibrants in our set of nucleosome standards: one that is symmetric *cis*-bivalent and one that is *trans*-bivalent. The bivalency sequential ChIP can then be normalized to either one of these standards, and because these two cases represent the limits of pulldown avidity, normalization to these calibrants will define the theoretical “range” in which true bivalency HMD (i.e. the proportion of nucleosomes with some bivalent configuration) exists (Extended Data Fig. 1e). We note that, because the signal from calibration to these standards are scalar multiples of each other, we cannot uniquely distinguish these two configurations in the genome. Absent any prior information about the dominant configuration of bivalency, the proportion of bivalently modified nucleosomes at a given locus will exist in the range defined by calibration to symmetric *cis-* or *trans*-bivalent standards (Extended Data Fig. 2a).

In practice, there are a few reasons why this is not a major concern. First, there is no mass spectrometry evidence that H3K4me3 and H3K27me3 exist on the same histone tail, despite specific enrichment for these marks and sensitive detection limits^49,81^, suggesting that configurations other than *trans-*bivalency are at most, extremely minor in abundance. Second, the scarcity of these *cis*-tail modifications is consistent with the bioschemical literature prior to this work that suggests the biogenesis of these *cis*-tail modifications is enzymatically challenging due to antagonistic allosteric effects (see Supplementary Note 4). Third, even if symmetric *cis*- bivalency does exist at some loci, for the purposes of tracking changes in bivalency across differentiation, we can still observe an increase or decrease in bivalency by this calibration method; we simply cannot precisely discern whether the effect is driven by nucleosomes gaining/losing *trans*-bivalency, *cis*-bivalency, or some combination of the two. The overall amount of bivalency would still increase or decrease in all those scenarios, and so long as our choice of calibrant remains consistent, we can still measure that change regardless of the calibrant that we use for our normalization. Therefore, though we have generated datasets using both calibrants, we present analyses of our reICeChIP bivalency pulldowns calibrated to the trans-bivalent standards.

### Supplementary Note 2

Throughout this study, we have defined gene promoters to be the region from 0 to +400bp relative to the TSS, representing the +1 and +2 nucleosomes of each gene. These nucleosomes tend to be well-positioned^83^ and, accordingly, are most likely to provide us with adequate read depth to robustly quantify each histone modification. This definition is conservative; we find that H3K4me3 and bivalent domains, which tend to be peak-like, have a median breadth of 550bp at bivalent genes (Extended Data Fig. 2d).

The width of these domains raises an important point regarding the measurement of histone modification density as a continuous variable. At a given nucleosome in a single allele of a single cell, there are only three possible states for a histone modification: symmetric, asymmetric or not present. However, nucleosome readers do not typically bind only a single nucleosome at a single position; rather, the local density of the modification across multiple nucleosomes is crucial in localizing these effectors through multivalent avidity-based interactions^84–87^. Indeed, we find that the HMD across sequential nucleosomes relative to the TSS is well autocorrelated (Extended Data Fig. 2e). This means that the interpretation of the HMD across a multinucleosomal span becomes more nuanced; a given histone modification may exist at one or more of those nucleosomes. Accordingly, despite the fact that a single nucleosome is essentially ternary in whether it has a given histone modification or not (i.e. HMD of 0% or 100%), a region spanning multiple nucleosomes could have an intermediate HMD; it is this latter quantity that is most relevant for the biological function imparted to the nearby genomic regions, and this is the quantity we analyse through this work.

### Supplementary Note 3

For the datasets presented in this work, the vast majority of promoters have a histone modification density between 0-100%, representing the proportion of nucleosomes at those promoters with the modification of interest (Fig. 2c; Extended Data Fig. 3a). However, at some loci, the measured HMD exceeds 100%. There are several possible reasons for this.

The most important of these possibilities is low input depth. The ICeChIP datasets are normalized to the input read depth at every genomic interval to accommodate for differences in local nucleosome density when computing the HMD. However, this means that at regions that are relatively nucleosome-depleted, there will be few reads in the input, meaning that the denominator of the HMD computation is quite small (Methods). This increased Poisson noise in these regions of low input can result in inflated apparent HMD beyond the physical limit of 100%. To accommodate for this, we can compute 95% confidence intervals for the HMD of each modification at each genomic position, and these confidence intervals virtually always overlap the physically possible range of HMD values (e.g., Fig. 1c). In naïve mESCs, only 0.5% of the promoters have a bivalency HMD above 100%, and for the vast majority of these promoters (86.1%), the 95% confidence interval error estimate ranges below 100%. The fact the apparent bivalency HMD calibrated by trans-bivalent standards, is broadly constrained to less than 100% further supports the idea that this choice of calibrant is appropriate and not inflationary (Supplementary Note 1).

There are also several other possibilities that are more challenging to accommodate for. First, some regions of the genome are known to be more artefact-prone for sequencing and mapping^88^; if the IP sample is enriched for these sequences relative to the input, then that could be disproportionately represented in the IP and have an apparent HMD greater than 100%. Second, the antibodies themselves could skew the apparent HMD. If the antibody is capturing substantial off-target material, then that will result in systematic inflation of the IP, resulting in an inflated HMD. Though ICeChIP barcoded nucleosome standards can help monitor off-target pulldown of some nucleosome species, we can only measure the capture of the standards that we actually have spiked into the experiment. If we do not have nucleosome standards available for a potential off-target modification, then we cannot definitively state that the antibody is not capturing that material. In this context, that is likely most important for H3K27me3 pulldowns; though we cannot state this definitively due to the lack of H3K27me2 standards, it is plausible that we are pulling down some amount of H3K27me2 with these IPs, resulting in slightly inflated apparent H3K27me3 HMD. However, this may not be too problematic; H3K27me2 and H3K27me3 are thought to be recognized by many of the same proteins and to have highly similar functions^82^, so the conflation of the two – if present – likely does not pose a significant problem in ascribing biologic function.

On a related note, at some loci, the bivalency HMD goes below 0%. In naïve mESCs, 8.8% of the promoters have a bivalency HMD below 0%, yet for the vast majority of these promoters (90.8%), the 95% confidence interval error estimate ranges above zero. This is because we employ *in silico* signal-correction for the bivalency dataset to remove signal that is attributable to H3K9me3. In essence, we can measure the amount of H3K9me3 pulldown in our bivalency ICeChIP dataset due to nucleosome standards employed, and we can separately measure the H3K9me3 HMD by a highly specific IP. We can then a linear combination correction matrix to remove the signal that is attributable to directly measured H3K9me3 at these loci. This method can effectively reduce the impact of modest off-target binding H3K9me3, but at some loci, will result in a subzero apparent HMD due to random sampling of read depth in the two distinct pulldowns employed.

Finally, at some sets of gene promoters, the trans-bivalency HMD is shown to be greater than the H3K4me3 or H3K27me3 HMD. This apparent discrepancy has a few possible reasons. First, there is some nuance in the interpretation of HMD in the context of single-target ICeChIP and reICeChIP. A nucleosome has two copies of each of its core histone proteins, including histone H3. This means that there are two possible sites of modification on each nucleosome for for each individual modification; if only one of those sites is modified, then that corresponds to an HMD of 50% because only half the possible modification sites are actually modified. However, this is different for the trans-bivalency HMD; by definition, only one trans-bivalency modification pattern can exist on a given nucleosome at any given time. If two “trans-bivalent” modification patterns existed on the same nucleosome simultaneously, then both H3K4 and both H3K27 residues would be trimethylated – which is symmetric *cis*-bivalency. As such, if one H3K4 and one H3K27 residue are trimethylated, then 100% of the possible trans-bivalency configurations for the nucleosome of interest are satisfied, meaning that the trans-bivalency HMD will be 100%. However, in this case, the H3K4me3 and H3K27me3 HMDs will only be 50% because only half the modifiable residues are actually modified.

The other caveat is that symmetrically modified nucleosomes will be pulled down more efficiently than asymmetrically modified nucleosomes due to avidity effects, as can be seen in the pulldown of symmetric vs. asymmetric H3K4me3 and *cis*-bivalency vs. *trans*-bivalency (Extended Data Fig. 2), and observed previously^46^. This means that calibration to symmetric nucleosome standards will have a larger denominator in computation of HMD and thereby yield lower apparent HMDs; this can also contribute to the lower apparent HMD of H3K4me3 and H3K27me3 relative to trans-bivalency. Accommodating for this phenomenon would require detailed profiling of asymmetric H3K4me3 (which is currently difficult due to the low quality of H3K4me0 antibodies), asymmetric H3K27me3 (which is not currently possible), and distinguishing between trans-bivalency and cis-bivalency (which is also not currently possible). However, as noted in Supplementary Note 1, so long as the method of calibration remains consistent, increases in apparent HMD will still correspond to increases in the modification of interest. Whether that increase in the target modification is due to asymmetric modification becoming symmetric or due to new gain of the modification at a previously unmodified locus in an instantaneous subpopulation remains unclear, but in both cases, modification density is still being gained at that locus. As such, even with these caveats, we can still quantitatively compare different datasets to each other as we use consistent calibration standards.

### Supplementary Note 4

Intriguingly, the catalytic activity of the EZH2-PRC2 core complex on nucleosome substrates is potentiated by pre-existing H3K27me3^89,90^, yet inhibited by H3K4me3, particularly when symmetric^49,56,59^. Conversely, symmetric H3K27me3 has been reported to modestly inhibit several of the human COMPASS-family complexes by qualitative assays, although only SET1 complexes were examined at the nucleosome level^60^. This presents a potential concern for our data – if the enzyme complexes that install these marks are mutually antagonized by the opposing mark, how might the widespread bivalency we observe arise? As the PRC2 effects are well established with detailed quantitative enzymology^49,56,59^, which we recapitulate (data not shown), we deployed more quantitative HMTase assays with a larger panel of relevant nucleosomal substrates to evaluate the COMPASS/SET1B/MLL-family core complexes for allosteric modulation by preexisting marks (Extended Data Fig. 6).

### Supplementary Note 5

In this context, sensitivity refers to the proportion of DEGs that are represented in a specific class of genes (e.g. H3K27me3-dominant bivalent genes), whereas specificity refers to the proportion of that class of genes that are differentially expressed. Under the prevailing bivalency model, bivalency is associated with poised genes that become upregulated or downregulated upon differentiation; as such, it should have high specificity for DEGs.

### Supplementary Note 6

The first way we evaluate different models for predicting DEGs is to compute the Bayes Information Criterion (BIC). Though not definitive, this metric estimates whether addition of a parameter to a model improves it more than would be expected from chance alone. When comparing two models, the model with the lower BIC will tend to have more explanatory parameters and/or fewer non-explanatory parameters than the model with the higher BIC. To this end, if BIC increases when a parameter is added, then it can be interpreted that the parameter being added contributes minimal additional explanatory power. Here, we find that adding bivalency to a model increases the BIC, meaning that it is likely (though not definitively) not contributing meaningfully more information in predicting DEG status in this differentiation paradigm.

A more definitive way to evaluate model accuracy is to use hold-out cross-validation. In this method, we split the set of all genes into two groups, one with 80% of the genes (the training set) and one with 20% of the genes (the testing set). We then train our GLMs on the training set and use the derived models to predict DEG status in the testing set. Hold-out cross-validation is a highly effective way of testing whether a model is overfit or underfit upon addition or removal of a parameter. If model accuracy increases substantially, then that would suggest the parameter has explanatory power over that provided by the other parameters. Conversely, if model accuracy decreases substantially, then that suggests that the additional parameter causes overfitting. Minimal changes in model accuracy suggest that the additional parameter contributes little to the model over the existing parameters, positively or negatively.

There are two metrics we use to test the accuracy of the predictions in the testing set. The first is by logistic regression thresholding, in which the gene is predicted to be a DEG if the modelled probability is greater than 0.5. The second is by computing the area under the receiver operator characteristic curve to measure true and false positive rates using different modelled probabilities as the thresholds.

Overall, we find that the GLM with bivalency barely changes model accuracy by either metric on hold-out cross-validation, with the magnitude of change being similar to that observed by instead adding H3K9me3 or DNA methylation. As such, we can interpret that none of these parameters – including bivalency – meaningfully contributes to the prediction of DEGs beyond what can be achieved with H3K4me3 and H3K27me3 in this system.

## Methods

### Cell Culture

Naïve mouse Embryonic Stem Cells (mESCs) were grown from the mESC E14 cell line (129/Ola background)^91^ in high glucose DMEM (Invitrogen), supplemented with 15%(v/v) FBS (Gibco), 1%(v/v) non-essential amino acids (Gibco), 1x penicillin/streptomycin (Gibco), 0.1 mM 2-mercaptoethanol (Gibco), 2mM L-glutamine (Gibco), 1000 U/mL LIF (ESG1107 Millipore), 3 µM CHIR99021 (LC Laboratories), 1 µM PD0325901 (LC Laboratories), sterilized using 0.1 µm filter flask (Millipore), stored up to 1 week in 4°C.

Primed mESCs were grown from the mESC E14 cell line (129/Ola background)^91^ in high glucose DMEM (Invitrogen), supplemented with 15%(v/v) FBS (Gibco), 1%(v/v) non-essential amino acids (Gibco), 1x penicillin/streptomycin (Gibco), 0.1 mM 2-mercaptoethanol (Gibco), 2mM L-glutamine (Gibco), 1000 U/mL LIF (ESG1107 Millipore), sterilized using 0.1 µm filter flask (Millipore), stored up to 1 week in 4°C.

Naïve and primed mESCs were grown on plates coated with 0.1% bovine gelatin (Sigma), grown to 70-90% confluence and passaged daily at a 1:3 ratio, with a media change 3 hours before passaging, supplemented with 1 vol. of fresh media 8 hours after passaging.

To initiate the adherent monolayer differentiation process to neuronal progenitor cells (NPCs; Day 0)^92,93^, naïve mESCs cells were split onto a gelatinized 6 cm plate at 1 x 10^4^ cells/cm^2^ and allowed to grow for 24 hours. On Day 1, the media was switched to RHB-A (Takara, Y40001) and was subsequently changed every other day. On day 4, cells were split and plated onto Poly-L-Ornithine, laminin-treated 6-cm plates. Prior to cell seeding the plates were treated with 0.01% Poly-L-Ornithine (Millipore, A004C) for at least 20 min, followed by 5 ug/cm^2^ of laminin (Fisher, CB40232) resuspended in basal RHB-A medium (Takara, Y40000). After washing off this treatment, cells were seeded in fresh RHB-A, supplemented with 10 ng/mL of bFGF (PeproTech, 100-18B) and EGF (PeproTech, 315-09). Cells were then split every 4 days at ≥ 20,000 cells/cm2 until an appropriate amount of NPCs were cultured for ICeChIP.

### Semi-synthetic Histone Preparation

Human histones H3.2(C110A)K4me3, H3.2(C110A)K9me3, H3.2(C110A)K27me3, H3.2(C110A)K4me3-K27me3 were made by semi-synthesis as described previously^46,94^. Asymmetric disulfide linked histone H3K4me3 - H3K27me3 dimers were made by semi-synthesis as described previously^56^.

### Octamer Reconstitution

Symmetrical H3K4me3, H3K27me3, H3K4me3-K27me3 and H3K9me3 octamers were made as previously described^95,96^. Briefly, equimolar amounts of histone H2A, H2B, H3 and H4 were mixed to the final concentration of 1 mg/ml in unfolding buffer (50 mM Tris-HCl pH 8, 6.3 M Guanidine-HCl, 10 mM 2-mercaptoethanol, 4 mM EDTA), subsequently they were loaded into 3500 M.W.C.O. dialysis tubing (Pierce Snakeskin) and dialyzed in 1000 volumes of refolding buffer (20 mM Tris-HCl pH 7.5, 2 M NaCl, 1mM EDTA, 5 mM DTT), overnight at 4°C. Dialyzed sample was 0.22 µm filtered, and octamers were resolved by S200 gel filtration chromatography (Superdex 200 10/300 GL, GE Healthcare) using refolding buffer as mobile phase. Eluted octamer fractions were pooled and concentrated using centrifugal filters (Amicon Ultra-4, 10k M.W.C.O., Millipore) to a final concentration of 5-15 µM, diluted with 1 volume of octamer storage buffer (20 mM Tris-HCl pH 7.5, 2 M NaCl, 1 mM EDTA, 5 mM DTT, 55% glycerol), and stored in −20°C. Concentration of octamer was measured spectroscopically using concentrator flow-through as a blank, ε_280nm_=44700 M^−1^cm^−1^, M_oct_≈108500 g^1^mol^−1^. Octamers were visualized using 18% separating (4% stacking) discontinuous Laemmli SDS-PAGE in Mini-Protean gel running system (Bio-Rad) run for 70 minutes at 22mA, 200V max.

Asymmetrical H3K4me3, H3K27me3 octamers were done as above with the following differences. Equimolar amounts of histone H2A, H2B, H3 and H4 were mixed in unfolding buffer to the total of 1-2 mg, where 90% of histone H3 was trimethylated on Lys 4 or Lys 27 and remaining 10% were unmethylated and had His_6_-tag at N-terminus with TEV cleavage site. Octamers were reconstituted overnight by dialysis in 1000 volumes of phosphate refolding buffer (50 mM Na_x_PO_4_ pH 7.5, 2M NaCl), at 4°C. Octamers were purified by S200 gel filtration chromatography, and his-tagged octamers were isolated using cobalt-based immobilized metal affinity chromatography Dynabeads magnetic particles. Octamers were incubated with magnetic beads for 10 min at 4°C on rotator, followed by two 1 ml washes (50 mM Na_x_PO_4_ pH 7.5, 2 M NaCl, 10 mM imidazole), and eluted with 50 ul of elution buffer (50 mM Na_x_PO_4_ pH 7.5, 2 M NaCl, 250 mM imidazole, 1 mM EDTA, 1 mM DTT), the elution step was repeated 6 times, fractions were characterized spectroscopically, pooled, diluted with 1 volume of octamer storage buffer, and stored in −20°C.

Asymmetrical *trans*-bivalent H3K4me3-K27me3 octamers were prepared the same way as symmetrical octamers with the following differences. Histones H2A, H2B, H4 and asymmetric disulfide linked histones H3K4me3 - H3K27me3 were mixed 1.2 : 1.2 : 1 : 0.5 ratio. Remaining steps were done as previously described but no reducing agents were used until octamer particles were formed.

All other octamers were obtained from EpiCypher, Inc.

### Nucleosome reconstitution

DNA barcodes were constructed based on 601 nucleosome positioning sequence^97^. One or both ends of 601 Widom sequence were substituted with 24 bp “barcode” sequence. Each barcode sequence is comprised of two 11 bp sequences absent in human and mouse genome, and constant 2bp linker DNA is added on a free end of the 601 nucleosome positioning sequence.

Nucleosomes were reconstituted as previously described^46^. Briefly, 10-100 pmol DNA and histone octamers were mixed in 1 : 1 ratio, at a final concentration >1 µM, and dialyzed in dialysis buttons (Hampton Research) against a non-linear gradient of sodium chloride 2M NaCl → 0.2M NaCl in a buffer containing 20 mM Tris-HCl pH 7.5, 1 mM EDTA, 10 mM 2-mercaptoethanol over the course of 12-16 hours^84^. Afterwards nucleosomes were recovered, diluted with 1 volume of 2x storage buffer (20 mM Na•Cacodylate pH 7.5, 10% v/v glycerol, 1 mM EDTA, 1x RL Protease Inhibitor Cocktail [1 mM PMSF, 1 mM ABESF, 0.8 μM aprotinin, 20 μM leupeptin, 15 μM pepstatin A, 40 μM bestatin, 15 μM E-64]), and stored at −20°C. Nucleosome concentration was measured by densitometry of 2% agarose gels, 1x TBE (89 mM tris-base, 89 mM boric acid, 2 mM EDTA) run for 30 minutes in 5V/cm electrical field gradient, followed by staining with 1x SYBR Gold (Invitrogen) for >30 minutes. Prior to electrophoresis, nucleosomes were disassembled with 2 M NaCl, roughly 1 pmol of nucleosomes were loaded per well and measured in triplicate against known quantity of free DNA of the same size. For use as ICeChIP standards, the semi-synthetic nucleosomes were diluted to 1 nM concentration using long-term storage buffer (10 mM Na•Cacodylate pH 7.5, 100 mM NaCl, 50% Glycerol, 1 mM EDTA, 1x RL Protease Inhibitor Cocktail, 100 µg/mL BSA(NEB)) and stored at −20°C.

### ICeChIP – input preparation

ICeChIP has been done as previously described^46,54^. Briefly, 10^7^-10^8^ plate adherent cells were released using Accutase (Millipore), quenched with complete medium and collected (500 rcf, 5 min., 4°C). Subsequent steps have been done on ice, cells after each wash were collected by centrifugation (500 rcf, 5 min., 4°C). Cells were washed twice with 10 ml PBS, twice with 5ml buffer N (15 mM Tris pH 7.5, 15 mM NaCl, 60 mM KCl, 8.5% (w/v) Sucrose, 5 mM MgCl_2_, 1 mM CaCl_2_ 1 mM DTT, 1x RL Protease Inhibitor Cocktail). Cells’ membranes was lysed by adding 1 volume of the 2x Lysis Buffer (Buffer N supplemented with 0.6% NP-40 substitute (Sigma)) to the single cell suspension resuspended in 2 PCVs (packed cell volumes) of Buffer N. After 10 minutes incubation on ice, nuclei were collected by centrifugation and resuspended in at least 6 PNV (packed nuclei volumes) of buffer N. Subsequently, nuclei were layered over 7.5 ml Sucrose Cushion N (15 mM Tris pH 7.5, 15 mM NaCl, 60 mM KCl, 30% (w/v) Sucrose, 5 mM MgCl_2_, 1 mM CaCl_2_ 1 mM DTT, 1x RL Protease Inhibitor Cocktail, 50 µg/mL BSA(NEB)) in a 50 ml centrifuge tube. Nuclei were spun through the sucrose cushion in swinging bucket rotor at 500 rcf for 12 min., 4°C. Nuclei were resuspended in 2 PNVs of buffer N, total nuclei acid content was measured spectroscopically at 260 nm by Nanodrop (Thermo Scientific) (1A_260_ = 50 ng/µl), prior to measurement, DNA was stripped from chromatin by adding 18 μl – 98 μl of 2 M NaCl and DNA was fragmented by vortexing and water bath sonication. The quality and quantity of nuclei was measured using a hemocytometer. The apparent concentration of chromatin was adjusted to 1 μg/μl with Buffer N. The following semi-synthetic standards were then spiked in: symmetrical H3K4me3, H3K9me3, H3K27me3, H3K36me3, H3K79me2, H3K4me3-K27me3 *cis*-bivalent, asymmetrical H3K4me3, H3K4me3-K27me3 *trans*-bivalent. To fragment the DNA, we aliquoted 100 µg of chromatin and added 1 Worthington unit of Micrococcal nuclease (Worthington) per 4.785 µg of chromatin (measured at 260nm, 1A_260_ = 50 ng/µl) and incubated at 37°C for 12 minutes. Digestion was stopped by adding 1/10 volume of 11x MNase stop buffer (110 mM EGTA, 110 mM EDTA pH 8.0). Subsequently, nuclei were lysed by slowly adding 5 M NaCl while mixing on a vortex (lowest setting) to the final concentration of 600 mM NaCl. Insoluble material has been spun down at 18000 rcf, 1min., 4°C.

Hydroxyapatite (HAP) chromatography has been done as described previously^46^. Briefly, 66 mg of HAP resin (Bio-Rad Macro-Prep^®^ Ceramic Hydroxyapatite Type I 20 μm) was rehydrated with 200 μl of HAP buffer 1 (3.42 mM Na_2_HPO_4_ and 1.58 mM NaH_2_PO_4_ final pH 7.2, 600 mM NaCl, 1 mM EDTA, 200 μM PMSF). Subsequently, 100 µg of digested soluble chromatin was added to the rehydrated resin and incubated for 10 minutes at 4°C on a rotator. Afterwards, resin slurry was transferred to the centrifugal filter unit (Millipore Ultrafree MC–HV Centrifugal Filter 0.45 μm). Resin was washed 4 times with 200 μl of HAP buffer 1, 4 times with 200 μl of HAP buffer 2 (3.42 mM Na_2_HPO_4_ and 1.58 mM NaH_2_PO_4_ final pH 7.2, 100 mM NaCl, 1 mM EDTA, 200 μM PMSF), and eluted 3 times with 50 µl HAP elution buffer (342 mM Na_2_HPO_4_ and 158 mM NaH_2_PO_4_ final pH 7.2, 100 mM NaCl, 1 mM EDTA, 200 μM PMSF), each wash/elution step was accompanied by centrifugation step (600 rcf, 30 sec., 4°C). Concentration of the chromatin was evaluated spectroscopically by Nanodrop (Thermo Scientific) (1A_260_ = 50 ng/µl). Apparent concentration of the chromatin was adjusted to 20 ng/µl with ChIP buffer 1 with BSA (25 mM Tris pH 7.5, 5 mM MgCl_2_, 100 mM KCl, 10% (v/v) glycerol, 0.1% (v/v) NP-40 substitute, 100 µg/ml BSA(NEB)).

### ICeChIP – immunoprecipitation

ICeChIP was performed as previously described^46^ with following modifications: α-H3K4me3 ChIP-5 µg chromatin, 2 µg 304M3B-1xHRV3C^53^, 40µl Streptavidin M-280 Dynabeads (Invitrogen). α-H3K9me3 ChIP-3 µg chromatin, 0.5 µg 309M3B^53^, 10 µl Streptavidin M-280 Dynabeads (Invitrogen). α-H3K27me3 ChIP - 0.8 µg chromatin, 0.6 µg CST C36B11 lot 8, 5 µl Protein G Dynabeads (Invitrogen). Aforementioned volumes of magnetic beads were washed twice with 200 µl ChIP buffer 1 with BSA. Antibodies were resuspended in 100 µl of ChIP buffer 1 with BSA; subsequently, magnetic beads were collected using magnetic rack, supernatant was removed, and magnetic beads were resuspended with the antibody solution and incubated for at least 1 hr, 4°C, on a rotator. Afterwards unbound antibody was washed away with two 200 µl washes of ChIP buffer 1 with BSA. Streptavidin beads were additionally washed twice with 200 µl of ChIP buffer 1 with BSA supplemented with 5 µM biotin for 10 min, at 4°C, on a rotator for each wash, followed by wash with 200 µl ChIP buffer 1 with BSA. Supernatant was removed on a magnetic rack and specific amount of chromatin, mentioned at the beginning of this chapter, was used to resuspend the magnetic beads. Chromatin was incubated with antibody-beads conjugates for 10-15 minutes, at 4°C, on a rotator.

Subsequently, magnetic beads were washed two times with 200 µl ChIP buffer 2 (25 mM Tris pH 7.5, 5 mM MgCl_2_, 300 mM KCl, 10% (v/v) glycerol, 0.1% (v/v) NP-40 substitute, 100 µg/ml BSA(NEB)), and one time with 200 µl ChIP buffer 3 (10 mM Tris pH 7.5, 250 mM LiCl, 1 mM EDTA, 0.5% Na•Deoxycholate, 0.5%(v/v) NP-40 substitute, 100 µg/ml BSA(NEB)), 10 minutes, at 4°C, on a rotator with tube change after each wash. These washes were followed by quick 200 µl ChIP buffer 1 (without BSA) wash, and 200µl TE wash (10 mM Tris-HCl pH 8.0, 1 mM EDTA). Chromatin was released from the resin using 50 µl ChIP elution buffer (50 mM Tris pH 7.5, 1 mM EDTA, 1% (w/v) SDS) at 55°C, for 5 minutes. Elution was supplemented with 200 mM NaCl, 10 mM EDTA, 10 µg of Proteinase K (Roche) and incubated at 55°C for 2 hours. DNA was isolated using 3 volumes of Serapure HD (1 mg/ml of 1 µm, hydrophobic, carboxylated, Sera-Mag SpeedBeads (GE 65152105050250), 20% PEG-8000, 2.5 M NaCl, 10 mM Tris pH 7.5, 1 mM EDTA, 0.05% Tween-20, filter sterilized prior to addition of magnetic beads), incubated for 5 minutes at room temperature, collected with magnetic rack and washed twice with >200 µl of 75% ethanol, on a magnetic rack without disturbing the magnetic beads. Subsequently, ethanol was carefully removed, and DNA was eluted by resuspending beads in 50 µl of TE buffer, magnetic beads were collected using magnetic rack and supernatant was moved to a new tube. In order to limit DNA loss, all operations have been performed using 250 µl siliconized tubes.

### reICeChIP

reICeChIP was performed in the same manner as described above with following changes. For the primary IP we have used 5 µg of HAP purified chromatin, 2 µg 304M3B-1xHRV3C – HRV 3C cleavable, biotinylated, αH3K4me3 Fab (PDB:4YHZ)^53^, immobilized on 40 µl Streptavidin M-280 Dynabeads (Invitrogen). After 10 minutes incubation of chromatin with antibody-resin conjugate, resin was washed three times with 200 µl ChIP buffer 1 with 100 µg/ml BSA, each wash consisted of 10 minutes incubation at 4°C, on a rotator, followed by tube change. Subsequently, resin was briefly washed with 200 μl ChIP buffer 1 with 100µg/ml BSA, and chromatin was released from the resin with 20 µl of ChIP buffer 1 supplemented with 100 µg/ml BSA and 4 µg HRV3C incubated (GE Healthcare) on ice for 60 minutes, elution step was repeated one more time and both elutions were combined. HRV 3C endoprotease efficiently cleaves at its target sites at 4°C^55^ and has a wide range of chemical tolerance^98^, but it highly-specific for its cognate cleavage sequence^99^ permitting facile use in ChIP under conditions which preserve nucleosomes. Primary elution was added to 0.6 µg CST C36B11, αH3K27me3 mAb, immobilized on 5 µl of Protein G Dynabeads(Invitrogen) for 10 minutes, at 4°C, on a rotator. Subsequently, magnetic beads were washed two times with 200 µl ChIP buffer 2, and one time with 200 µl ChIP buffer 3, 10 minutes, at 4°C, on a rotator, with tube change after each wash. These washes were followed by quick wash with 200 µl ChIP buffer 1 (without BSA). Chromatin was released from the resin by 5 minutes incubation with 50 µl ChIP elution buffer, at 55°C. Chromatin was Proteinase K digested, and DNA was purified as described in ICeChIP – immunoprecipitation chapter.

### Design, expression, and purification of 304M3B-1xHRV3C

304M3B-1xHRV3C Fab is based on previously described Fab 304M3B(PDB:4YHZ)^53^. The gene encoding the Fab was modified to contain HRV3C cleavage site at the C-terminus of the heavy chain. To that end, we inserted SSSLEVLFQGP (AGC AGC AGC CTT GAA GTC CTC TTT CAG GGA CCC) sequence just after the position T229 of heavy chain (numbered as in PDB:4YHZ) and before biotinylation acceptor peptide (GLNDIFEAQKIEWHE)^100^. The Fab was expressed in the 55244 strain of *E.coli* in the TBG media (Terrific Broth (FisherBrand), 0.8% (v/v) glycerol) with 100 μg/ml carbenicilin, grown for 24 hours, at 30°C, 200 rpm in the Fernbach non-baffled flasks, with constricted airflow. Fab was purified using Protein G-A1^101^ affinity chromatography, followed by cation-exchange chromatography (Resource S, GE Healthcare). Purified Fab was *in vitro* biotinylated using BirA biotin ligase.

### Sequencing and Data Analysis

Each sequencing library was made using 10 ng of DNA. Illumina sequencing libraries were made with NEBNext Ultra II DNA Library Prep for Illumina (NEB), according to the manufacturer protocol. DNA libraries were amplified using 8 PCR amplification cycles (C1000, Bio-Rad). Cluster generation and sequencing was performed using the standard Illumina protocols for Illumina HiSeq 4000 by the University of Chicago Functional Genomics Core facility. Data analysis was performed as previously described^46^. Briefly, reference genome was modified to contain sequences of semi-synthetic nucleosome barcodes. Reads were mapped to GRCm38/mm10+barcodes reference genome, using Bowtie2^102^, end-to-end, sensitive preset. Subsequently, SAM files were filtered to reject unmapped and unpaired reads, as well as fragments with length > 200bp and Phred quality score < 20. Paired reads were merged into single interval of the fragment. BEDTools^103^ were used to calculate bedgraphs of the genome coverage. IP and input bedgraphs of genome coverage were subsequently merged into multiple entry interval file of genome coverage. Number of DNA fragments coming from semi-synthetic nucleosome standards were counted for each barcode and IP efficiency was calculated for each histone mark.

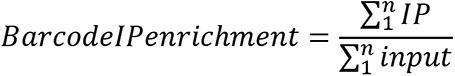

where, n is the identity of the n-th barcode assigned to the specific histone mark.

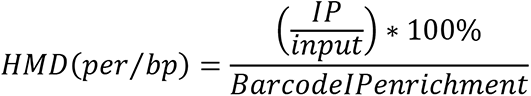

where, IP and input refer to the depth of the genome coverage at any given position of IP and input for specific histone mark. An IP efficiency can be interpreted as maximal yield of the IP for a given histone mark, or 100% HMD. However, there is a number of factors that can lead to apparent HMD values greater than 100% including: uncertainty due to random sampling of sequenced fragments, in that case standard deviation is approximately equal to square root of sequencing depth, or off-target capture by antibody, where the off-target histone mark or combination of histone marks have greater IP efficiency than the antibody intended target mark.

For all analyses, the HMD averaged over the N+1 and N+2 nucleosomes (taken to be 0 to +400bp into the gene body) was employed as representative of the promoter—this captures the most substantial H3K4me3 and H3K27me3 enrichment.

Genomic browser views were made using IGV. Heatmaps and gene ontology analysis was made using Homer Software^104^. Further analysis and sectioning of data was conducted in R using the R code provided in Data Availability. Plots were made using ggplot2 and Microsoft Excel. For the bivalency tracks, *in silico* antibody off-specificity correction was performed as previously described for H3K9me3^46^. ICeChIP-qPCR was performed with previously described primers^46^.

### Analysis of External Data

Bisulfite sequencing data was obtained from GEO series accession number GSE41923, dataset accession numbers GSM1027571, GSM1027572, GSM1027573, and GSM1027574. Methylation count files were obtained for each dataset and lifted to mm10. The average methylation for each promoter was then calculated for the 0 to +400bp region relative to the TSS of Refseq promoters using BEDTools.

Bulk RNA-seq data was obtained from GEO series accession numbers GSE108832 and GSE65697, dataset accession numbers GSM2913929, GSM2913930, GSM2913931, GSM1603282, GSM1603283, GSM1603284, GSM1603285, GSM1603286, and GSM1603287. Pseudoalignment was conducted against the Refseq mm10 transcriptome using kallisto^105^ with fragment length mean and standard deviation of 200 and 20, respectively, and 100 iterations. Pseudoalignments were then loaded into R for differential expression analysis using sleuth^106^, with correction for batch effects between primed mESCs and NPCs due to contribution to principal components of the same. Differentially expressed genes were identified as *q ≤ 0.05*. Single-cell RNA-seq data was obtained from GEO series accession number GSE113417 and aligned as above with kallisto.

Suz12 ChIP data to measure PCR2 localization for WT, Ezh2 KO, and Ezh1 KO/Ezh2 KO cells was obtained from GEO series accession number GSE116603, dataset accession numbers GSM3243624, GSM3243625, and GSM3243626. Peak files were obtained for all these datasets lifted to mm10. Ezh2 peaks were identified as peaks lost in Ezh2 KO relative to WT cells. Ezh1 peaks were identified as peaks lost in Ezh1 KO/Ezh2 KO relative to Ezh2 KO cells.

Set1A ChIP data was obtained from GEO series accession number GSE98988, dataset accession numbers GSM2629676, GSM2629677, GSM2629678, and GSM2629691. FastQ files were downloaded for the input and ChIP datasets for each replicate, then aligned to mm10 using Bowtie2 in end-to-end mode with the sensitive preset. Peak calling was then conducted on the alignments with MACS2^107^, and consensus peaks for each replicate were identified.

Mll2 ChIP data was obtained from GEO series accession number GSE78708, dataset accession number GSM2073022. Peaks were obtained and lifted to mm10.

### Methyltransferase assays

Enzymatic complexes were procured from Reaction Biology Corporation. Methyltransferase reactions were done using following concentrations of the enzymatic complexes: 200 nM hsMLL1(3745-3969), 200 nM hsMLL2(5319-5537), 400 nM hsMLL3(4689-4911), 200 nM hsMLL4(2490-2715), 800 nM hsSet1A(1418-1707), 800 nM hsSet1B(1629-1923), in a complex with hsWDR5(22-334), haRbBP5(1-538), hsAsh2L(2-534), 2x(hsDPY-30(1-99)), supplemented with 4%(v/v) RBC MLL enhancer (Reaction Biology Corp); 800 nM hsEzh1(2-747), 120 nM hsEzh2(2-746), in a complex with hsAEBP2(2-517), hsEED(2-441), hsRbAp48(2-425) and hsSUZ12(2-739) supplemented with 3.6mM hsJarid2(119-574) provided by Dr. Peter Lewis’s laboratory. 30 ng/µl of semi-synthetic nucleosome substrate, 10 µM [^3^H]-SAM (50-80 Ci/mmol, Perkin Elmer Health Sciences), and enzymatic complexes were mixed in the Reaction Buffer (50 mM Tris pH 8.0, 91 mM NaCl, 5 mM MgCl_2_, 1 mM DTT, 10% glycerol, 1 mM PMSF) and incubated at 30°C. At designated time points, 4 μl of reactions were spotted on P81 Ion Exchange Cellulose Chromatography Paper (Reaction Biology Corp). Spotted paper was washed 4 times with 250 ml of 50 mM NaHCO_3_ pH 9.0, for 5 minutes on a platform shaker, briefly washed with acetone, air-dried and immersed in scintillation fluid. ^3^H decay rate was measured by scintillation counter (LS 6000IC, Beckman).

## Data and Software Availability

ICeChIP-seq data generated for this study has been deposited at the Gene Expression Omnibus (GEO) under accession numbers GSE108747 and GSE183155. R markdown file for analysis and sectioning of datasets is provided at https://www.github.com/shah-rohan/bivalency/.

